# Sugar alternatives shift the evolutionary trajectory of fluoride resistance in cariogenic and non-cariogenic dental bacteria

**DOI:** 10.64898/2026.11.16.767098

**Authors:** Madeline R. Shay, Megan A. Eakin, Randy B. Stockbridge, Livia M. A. Tenuta, Xiaoxia Nina Lin, Kevin B. Wood, Luis Zaman

## Abstract

Dental caries result from acidogenic biofilms dissolving enamel while growing on fermentable dietary sugars. While fluoride from toothpastes and professional products is widely used for caries control, sugar alternatives with inhibitory effects on some dental microbiota (e.g., erythritol and xylitol) instead reduce the cariogenicity of foods and act as prebiotics, often in combination with of fluoride, to control caries progression. Despite their growing prevalence, we do not understand how frequent exposure to sugar alternatives impacts the evolutionary pressures faced by oral bacteria. Herein, we tested the effects of carbon source (glucose, sorbitol, erythritol, or xylitol) on the modes and effectiveness of evolving fluoride resistant cariogenic (*S. mutans*) and non-cariogenic (*S. gordonii)* populations. For *S. gordonii,* evolving in glucose resulted in significantly more non-synonymous mutations compared to populations evolving in sugar alternatives (erythritol, sorbitol, xylitol), but without a drastic increase in fluoride resistance. Contrastingly, for cariogenic *S. mutans,* evolving in the presence of fermentable sugars resulted in much greater fluoride resistance, but without higher numbers of substitutions relative to those found in isolates from populations evolved in the presence of non-fermentable sugars. We observed previously unreported fluoride resistance mutations across carbon conditions, including a case of parallel evolution in the histone-like binding protein (smu_589) in *S. mutans* isolates evolved in environments with fluoride and a variety of sugar alternatives. Together, these results demonstrate that commonly used sugar alternatives can shape the evolutionary trajectory of dental microbiota to a broad stressor, such as fluoride.

## Introduction

Early childhood caries (ECC) is a common infancy disease leading to pain, infection, missed school productivity, and significant healthcare costs^1^. ECC is linked to a dysbiosis in the dental biofilm, with an increased presence of cariogenic community members, such as *Streptococcus mutans* and *Candida albicans*, wherein frequent exposure to fermentable sugars creates an acidic environment favoring mineral loss from the tooth structure and caries lesions progression.

*S. mutans* is considered a core species in ECC^2^, as it is an early colonizer that can modify the biofilm matrix by secreting extracellular polysaccharide-producing enzymes. Once established in the biofilm, frequent consumption of fermentable sugars allows *S. mutans* to create a niche with enhanced selection for acidogenic and aciduric strains, accelerating caries lesions formation^3,4^. In contrast, *Streptococcus gordonii* is considered a commensal pioneer colonizer that can have beneficial effects on the dental microbiome. *S. gordonii* can directly colonize the acquired salivary pellicle, and competes with *S. mutans* for binding sites and nutrients, which can mitigate the growth of acidogenic strains and limit caries progression^5^.

Reducing the frequency of exposure to fermentable carbohydrates is among the first-line defense used to slow caries progression. Sugar alternatives (e.g. erythritol, xylitol, and sorbitol) are widely used in “sugar-free” food products and “cavity-friendly” candies due to their low caloric content, and intense sweetness^6–8^. In addition to their direct dietary effects, some sugar alternatives seem to have intrinsic anti-cariogenic properties and can inhibit the growth and biofilm formation of important dental microbiota, such as *S. mutans*^9,10^. Clinical isolates with resistance to xylitol, likely from repeated dietary exposure, have been observed to be less cariogenic than sensitive strains^11^.

Varying the carbon source used in laboratory grown dental biofilms can drive the metabolic state of individual community members, modulate cariogenic potential via acid production, and alter community composition. On one hand, some sugars like sucrose dramatically enhance biofilm production since it acts as a substrate for extracellular polysaccharide production. On the other, commonly used non-fermentable sugar alternatives such as xylitol and erythritol have been shown to have inhibitory effects on biofilm formation, further shifting inter-species interactions amongst cariogenic community members^9,10,12,13^. Consistent with these observations, products combining xylitol with fluoride in toothpastes can reduce demineralization and, at high xylitol concentrations, enhance enamel remineralization relative to fluoride-only products^14,15^.

Fluoride is the most commonly used oral agent to control caries development, acting primarily by reducing tooth demineralization and enhancing remineralization, with significant antimicrobial effects only apparent at high concentrations^16–20^. Exposure to fluoride can cause a drastic shift in the metabolic activity of dental microbiomes *in vitro*, resulting in reduced sugar metabolism within the community and changes to community composition^21^. At the individual level, fluoride can cause broad cellular stress^22–24^ and, in *S. mutans*, eventually leads to cell lysis via an unmitigated competence cascade^25^.

Herein, we investigated the effects of chronic exposure to sugar alternatives on the evolution of fluoride resistance in a representative cariogenic strain, *S. mutans*, and a representative non-cariogenic strain, *S. gordonii* (**Fig. 1**). Though the effects of several sugar substitutes on dental biofilm growth have been studied with synthetic consortia on ecological timescales, there are still open questions about the evolutionary consequences of prolonged exposure to sugar alternatives on cariogenic and commensal community members. We observed that *S. mutans* populations that evolved in the presence of non-fermentable sugar alternatives (erythritol or xylitol) acquired less fluoride resistance than those evolved in fermentable sugars (glucose or sorbitol). Furthermore, we identified several intergenic and genic regions commonly mutated in fluoride resistant isolates. Some of the observed mutations matched commonly mutated genes in fluoride resistant clinical isolates, while others were novel, such as a previously unreported mutation in a histone-like binding protein (smu_589, HLP). These results demonstrate that fluoride is a stressor that can drive adaptation and resistance in oral microbes, that commensal and pathogenic strains differentially evolve resistance, and the specific mechanisms of that resistance depend on the added sugar or sugar alternative.

**Figure 1:**
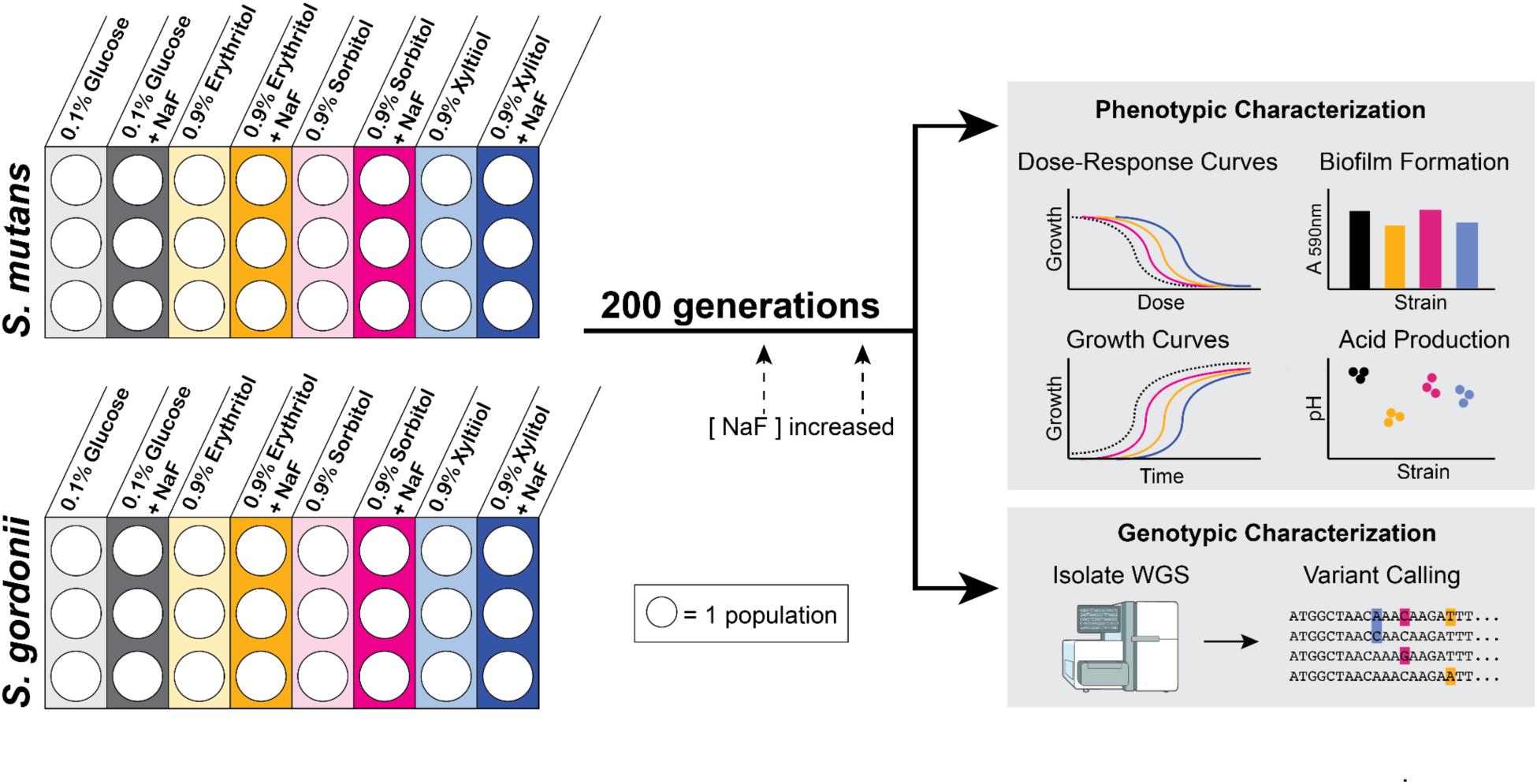
The workflow for the evolution experiment. Three replicate populations of each strain in each condition evolved for 200 generations. For the populations exposed to fluoride, the fluoride concentration was increased twice during the experiment according to the fitness of the population at that time. After 200 generations, dose response curves were determined for the bulk populations in response to fluoride and inhibitory sugar alternatives. Isolates were obtained from the bulk populations and analyzed via short read whole genome sequencing, tested for their fluoride sensitivity, and characterized using growth curves in various fluoride and carbon environments.

## Materials & Methods

### *In vitro* Evolution to Fluoride & Sugar Alternatives

*Streptococcus mutans* UA159 (ATCC #700610) and *Streptococcus gordonii* V288 (ATCC #35105) were serially passaged for 200 generations in TSB medium without dextrose (VWR Cat. # 90004-068) supplemented with 0.1% w/v glucose, 0.9% w/v erythritol (ThermoFisher Cat. # AAA1581322), 0.9% w/v sorbitol (Millipore Sigma Cat. # S1876-100G), or 0.9% w/v xylitol (Millipore Sigma Cat. # X3375-100G) in the presence or absence of sodium fluoride (ThermoFisher Cat. # S299-100) (**Fig. 1**). The number of generations was calculated according to the dilution factor and number of passages, and was consistent across carbon conditions. All carbon and fluoride supplements were prepared in a concentrated stock, filter sterilized, and diluted to their final working concentration. There were three replicate wells per condition. The concentration of the carbon supplements was held constant over the course of the experiment. For the populations exposed to fluoride, the concentrations were chosen according to the ancestor’s IC_50_ value for sodium fluoride in the respective carbon condition at 20 hrs; the concentration of sodium fluoride was increased according to the population’s IC_50_ at 150 and 175 generations (**Suppl. Fig. 1**). The experiment was done in 96 well plates (ThermoFisher Cat. #12-556-008) sealed with Breathe Easier films (Diversified Biotech Item #BERM-2000), static, under 5% CO_2_ at 37°C. Every 24 hours each well was mixed and diluted 1:25 in fresh medium, with OD_600nm_ measured at each transfer. After an aliquot was transferred to fresh medium, the 96 well plate from the previous passage was prepared for storage via the addition of glycerol (final concentration 30% v/v), sealed (Fisher Scientific Cat. #07-200-684), and stored at -80°C. An isolate from each bulk population was selected by streaking the frozen stock on TSB agar plates, randomly selecting a colony, and re-streaking on TSB agar. A single colony was grown in TSB with dextrose overnight, mixed with glycerol (30% v/v), and stored at -80°C for follow up assays. Sensitivity of the endpoint bulk populations and the respective randomly selected isolates to fluoride and the sugar alternatives (erythritol, sorbitol, xylitol) were quantified. Additionally, the maximum growth rate in the presence and absence of fluoride, the biofilm formation potential, and the acid production of the isolates was determined and compared to the ancestor. Lastly, these phenotypes were paired with whole genome sequencing of the randomly selected isolate from each population to identify the modes of adaptation.

### Isolate Whole Genome Sequencing

DNA was extracted from isolates using the GeneJet Genomic DNA Purification kit (ThermoFisher) and sent to SeqCoast Genomics (Portsmouth, NH, USA) for short-read whole genome sequencing with a 200 Mbp/1.3 Million read depth. Sequencing data was aligned to NCBI genome assembly ASM746v2 and ASM973053v1, for *S. mutans* and *S. gordonii* respectively, using breseq version 0.39.0 and Bowtie2 version 2.5.4^26^.

### Genetic Engineering

Information on the primers and constructs used for homologous recombination can be found in **Suppl. Table 1**. Regions with point mutations of interest were amplified from strains with the genotype using colony PCR Q5 Hot Start High-Fidelity 2X Master Mix (NEB Cat # M0494S). Then, amplified constructs were introduced to the ancestor strain of *S. mutans* according to published methods with minor modifications (**Suppl. Method 1**)^27^. Transformations were carried out in CDM with L-asparagine^28,29^, and under 5% CO_2_ at 37°C, and plated on TSB with dextrose agar. After plating, colonies were grown in TSB with dextrose for 24 hours under the same conditions, screened for the mutation using colony PCR, then re-streaked and screened until a pure mutant isolate was identified and confirmed via sanger sequencing (**Suppl. Table 1**).

### Fluoride Dose Response Curves

The fluoride resistance for all bulk populations and isolated strains was tested in TSB without dextrose supplemented with carbon and sodium fluoride according to each assay condition. Starter cultures were grown in TSB (ThermoFisher Cat. # DF0370-17-3) static under 5% CO_2_ at 37°C, and then diluted 1:100 into their respective assay conditions in 96-well plates. The 96-well plates were sealed with a Breathe Easier film and allowed to grow for 22-24 hours. At the endpoint, the wells were thoroughly mixed and OD_620nm_ was read using a Byonoy microplate reader. Data from each replicate (n=4 for isolate, n=3 for bulk populations) was fit to a Hill model (4-parameter logistic curve) relating the maximum growth (OD_620nm_) at the endpoint and the concentration of fluoride to determine an IC_50_ value (**Equation 1**), with any outliers identified and removed according to a two-sided Grubb’s test (α=0.05). U-M GPT was used to generate code for dose response curve data analysis and IC_50_ determination, all code was reviewed and cross-validated using a sub-set of the data analyzed using GraphPad Prism.

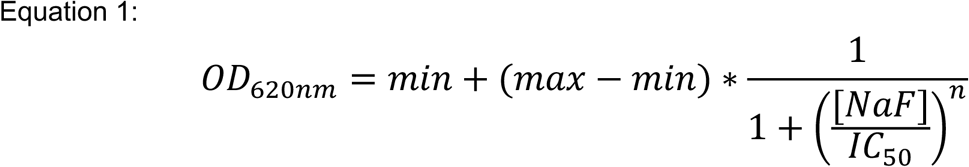

OD620nm= growth at endpoint (i.e., response)

min= minimum OD620nm

max= maximum OD620nm

[NaF]= concentration of fluoride

IC50= dose at which the OD620nm is 50% of the way between the min and max

n= Hill coefficient (i.e., steepness of the dose-response curve)

Remaining replicates were then combined per condition to determine mean inhibitory concentration 50 % (IC_50_) ± standard deviation (SD). A Welch’s t-test was used to compare the mean IC_50_ for each evolved population or isolate to the ancestral IC_50_, with the p-value reported in **Supporting Information**.

Grand averages of the fluoride IC_50_ of the bulk populations and isolates evolved in fermentable sugars (glucose and sorbitol) and fluoride were compared to those evolved in non-fermentable sugars (erythritol and xylitol) and fluoride using a one-way ANOVA with a Tukey’s Honest Significant Difference (HSD) test. These data are shown in **Fig. 2** with summary statistics in **Suppl. Tables 2&3**.

**Figure 2:**
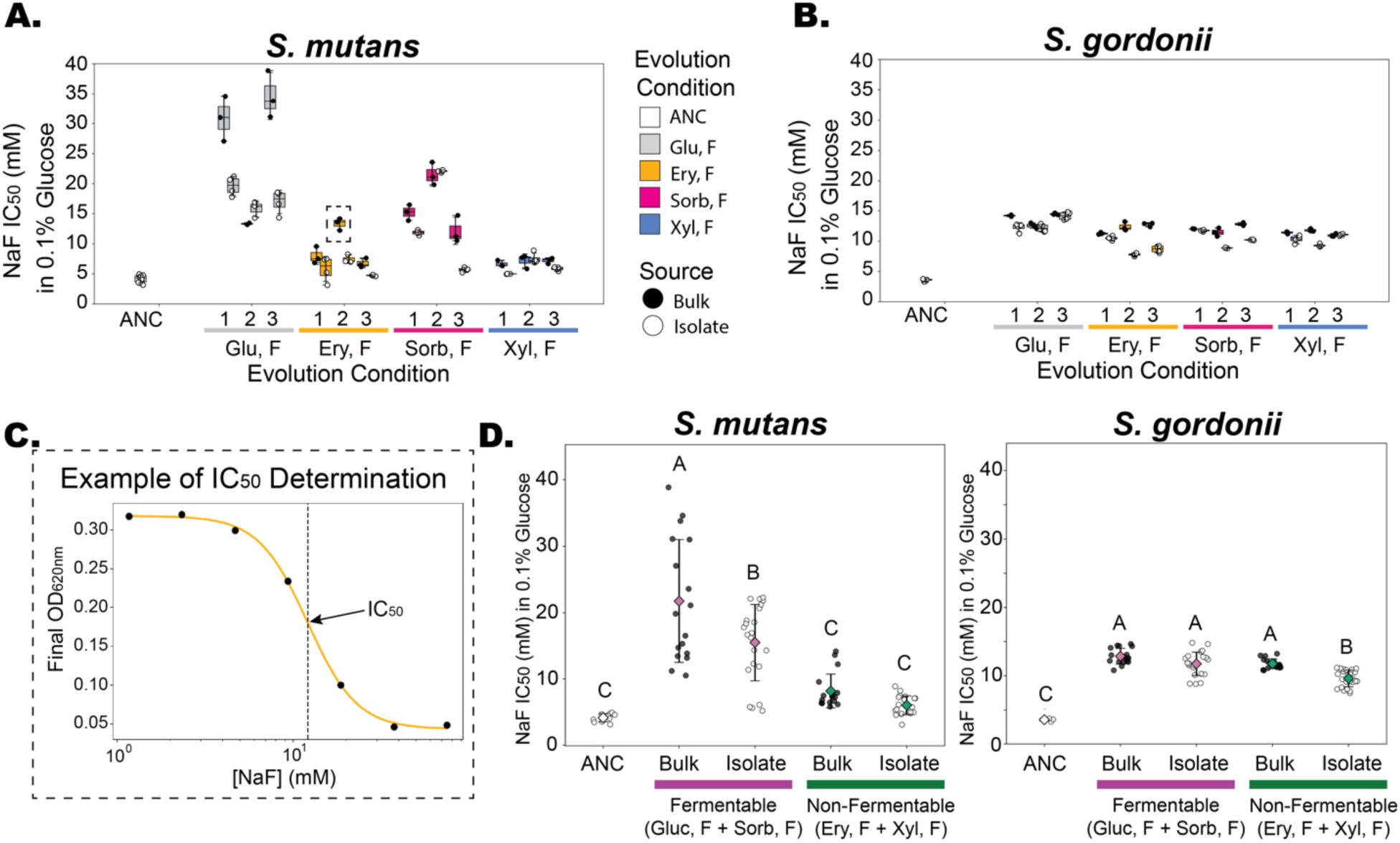
The fluoride IC_50_ in 0.1% glucose was determined for the bulk populations (●) and isolates (○) of (**A**) *S. mutans* and (**B**) *S. gordonii.* Evolution conditions are as follows: (ANC)= ancestor, (Gluc,F)= 0.1% w/v glucose and sodium fluoride, (Ery F)= 0.9% w/v erythritol and sodium fluoride, (Sorb F)= 0.9% sorbitol and sodium fluoride, (Xyl F)= 0.9% w/v xylitol and sodium fluoride. Fluoride IC_50_ of the evolved bulk populations and isolates at 200 generations was determined by fitting each four-parameter logistic dose response curves to 8 data points associating fluoride concentrations and final ODs, an example of the IC_50_ fit of the data shown in a boxed square in panel **A** is shown in panel **C**. To visualize the effects of fermentable (glucose & sorbitol) and non-fermentable (erythritol & xylitol) carbon source on fluoride resistance, the bulk populations and isolates were batched by the type of carbon condition they evolved in (**D**). A one-way ANOVA with Tukey HSD comparisons compared the fluoride IC_50_. Groups with the same letter indicate no statistically significant difference between groups, whereas groups with no shared letters are significantly different from each other based on the adjusted p-value (*α* = 0.05). Summary statistics are available in **Suppl. Tables 2 & 3**. Fluoride dose response data and fit IC_50_ values in other carbon conditions can be found in **Suppl. Fig. 4** and the **Supporting Information**.

### pH Determination

Turbid overnight cultures were diluted 1:100 in TSB without dextrose supplemented with 0.1% w/v glucose, transferred to a 96 well plate which was then sealed with a Breathe Easier film (Diversified Biotech Item #BERM-2000), and grown static for 24 hours at 37°C with 5% CO_2_. There were four replicates per strain. The samples were centrifuged to pellet the cells (4,000 rpm for 10 minutes), and 100μL of the supernatant was transferred to a new 96 well plate. 10μL of pH-indicator solution pH 4.0-10.0 (Millipore Sigma, Cat. #1.09175) was added to each well, and the absorbance spectra (300 nm - 700 nm) was measured for each sample on a Biotek Cytation5 (Aligent). A standard curve of calibrated fresh media was used to fit 4 parameter logistic curves comparing the absorbance at 520nm/430nm and 620nm/520nm (**Suppl. Fig. 2**). The 520nm/430nm absorbance ratio was used to determine the pH below 5.5, whereas the 620nm/520nm absorbance ratio was used to fit the pH above 5.5. A Welch’s t-test with a Holm adjustment (*α* = 0.05) was used to compare the pH of each isolate to that of the ancestor for that strain (**Supporting Information**).

### Biofilm Quantification

The biofilm-forming potential of isolates was tested in 96 well round bottom plates, with an n=4 per strain. Biofilm formation was tested in TSB without dextrose supplemented with 1% w/v sucrose, glucose, erythritol, sorbitol, or xylitol. Wells were inoculated 1:100 with turbid overnight culture, and grown static at 37°C with 5% CO_2_ for 24 hours. At the endpoint, crystal violet staining was done according to established methods ^30^. The solubilized crystal violet stain was diluted 4 X into 30% acetic acid so that the maximum absorbance at 590 nm was within the dynamic range of absorbance using a Biotek Cytation5 spectrophotometer (Aligent). A Welch’s t-test with a Holm adjustment (*α* = 0.05) was used to compare the pH of each isolate to that of the ancestor for that strain (**Supporting Information**).

### Kinetic Growth Curves

Growth curves were conducted in TSB media without dextrose supplemented with carbon (glucose, sorbitol, erythritol, or xylitol) and sodium fluoride according to each assay. Starter cultures were grown static under 5% CO_2_ at 37°C, and then diluted 1:100 into 96 well plates with the assay conditions, with 3 replicates per condition. The plates were sealed with Breathe Easy films (Diversified Biotech Item #BEM-1), and grown under 5% CO_2_ at 37°C, with OD_600nm_ measurements every 3 minutes with intermittent shaking over 24 hours using a Biotek Cytation5 (Aligent). The data were log-transformed and smoothed with a Savitzy-Golay filter (window size 21 minutes, polynomial order 2) for determination of the maximum specific growth rate (μ_max_) and the time of the maximum growth rate. The μ_max_ was the maximum derivative of ln (OD_600nm_) after the first hour of growth. All growth parameters were calculated per biological replicate, with the reported statistics being mean ± standard deviation (SD) for n=3. Plots showing the fit points for μ_max_ including replicate variability and additional curve fitting information for the evolved isolates in each carbon environment can be found in **Suppl. Fig. 3**.

### Sugar Alternative Dose Response Curves

The effect of sugar alternatives (erythritol, sorbitol, or xylitol) on the growth of the evolved bulk populations and isolates were tested with an 8-point dose response curve with 2 fold serial dilutions from 10% w/v to 0% (n=4). A high concentration stock of the sugar alternatives (20% w/v) was prepared via sterile filtration, serially diluted, and mixed 1:1 with 2X TSB without dextrose. Starter cultures were grown in TSB static under 5% CO_2_ at 37°C, and then diluted 1:100 into 96 well plates with the assay conditions. The 96 well plates were sealed with a Breathe Easier film and allowed to grow for 22-24 hours. At the endpoint, the wells were thoroughly mixed and OD_620nm_ was read using a Byonoy microplate reader.

### Area Under the Curve (AUC) Quantification & Analysis

The area under the curve (AUC) was calculated independently for each replicate (n=4). All replicates were baseline corrected by subtracting the media blank with any negative values set to zero. AUC was calculated using the trapezoidal rule according to **Equation 2**, wherein the AUC is the cumulative growth (OD_620nm_) over the dose range. The mean AUC and standard deviation were calculated for each bulk population or isolate (**Suppl. Fig. 5**). The mean AUC for each evolved sample was compared to that of the ancestor strain with a Welch’s t-test, with the t-statistic and p-value reported in **Supporting Information**.

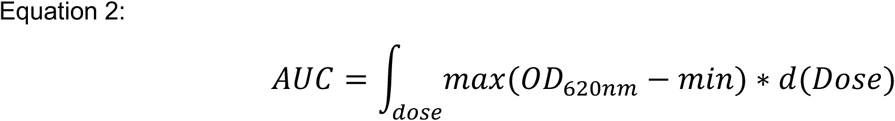

AUC= area under the curve

Response= growth (OD620nm)

min= OD620nm of media blank

d(Dose) = integration with respect to change in dose

Dose= sugar alternative concentration

## Results & Discussion

### Evolved Isolate & Bulk Population Fluoride Response

*S. mutans* and *S. gordonii* were evolved in the presence of glucose (0.1% w/v) or sugar alternatives (0.9% w/v erythritol, sorbitol, or xylitol) for 200 generations in the presence and absence of sodium fluoride (**Fig. 1**). If fluoride was present, it was sequentially increased according to the tolerance of each population at 150 and 175 generations (**Suppl. Fig. 1**).

To characterize the phenotypes of evolved populations, one isolate was randomly selected per replicate population after evolving in fluoride for 200 generations and was evaluated for fluoride resistance in varying carbon environments. Notably, for *S. gordonii,* carbon environment during the evolution experiment had a minimal effect on the degree of fluoride resistance in 0.1% w/v glucose; for *S. mutans* the presence of a fermentable sugar (glucose or sorbitol) in the evolution condition drastically increased the fluoride IC_50_ relative to the non-fermentable sugar alternatives (erythritol or xylitol) with a mean difference of 13.6 mM for the grand averages of the bulk populations (**Fig. 2D**). These findings indicate that non-fermentable sugars can alter the evolutionary trajectory of fluoride resistance in cariogenic *S. mutans* and, in this case, limit acquired fluoride resistance. In contrast, carbon source had little effect on the trajectory of fluoride resistance in the commensal *S. gordonii*, which suggests the effects of sugar alternative on adaptability to fluoride stress is species specific.

Fluoride resistance of the bulk populations was also tested and compared to the respective randomly selected isolate, wherein some isolates had significantly lower fluoride resistance than the bulk population they were selected from. When comparing the grand averages of the bulk populations and isolates evolved in fermentable sugars, there is a mean difference of 6.3 mM for *S. mutans* with a mean difference of only 1.1 mM in *S. gordonii* (**Fig. 2D, Suppl. Tables 2 & 3**). This discrepancy may indicate heterogeneity in some of the evolved populations, most commonly in *S. mutans* populations evolved in fermentable sugars (glucose, sorbitol). Heterogeneity in the evolved *S. mutans* populations could increase the resistance of the culture to fluoride treatment in several ways. Bulk populations could be harboring genetic variation on which selection could act, or they could be facilitating cooperative or competitive intra-population dynamics. Additionally, *S. mutans* growing in fermentable sugars would produce more acid, wherein the populations would experience more dynamic and drastic pH fluxes each passage relative to the populations growing on non-fermentable sugars. These more pH-variable environments may explain the seemingly species- and carbon-specific phenotypic heterogeneity in the bulk populations due to the proton coupling of the *S. mutans* fluoride efflux pump (smu_1289)^31^.

Fluoride resistance was also measured in environments containing the sugar alternative in which each bulk population or isolate had evolved (**Suppl. Fig. 4**). Fluoride resistance levels varied significantly between many of the bulk populations and isolates. Notably, one isolate evolved in erythritol and one isolate evolved in sorbitol showed no significant increase in fluoride resistance relative to the ancestor when tested in the sugar alternatives they evolved in, surprisingly losing the benefit we observed in glucose environments. These observations, combined with the discrepancies in the fluoride resistance when tested in glucose, are suggestive of heterogeneous populations balancing trade-offs between their adaptation to sugar alternatives and fluoride resistance. Consistent with such a trade-off, evolved fluoride resistance in both *S. mutans* and *S. gordonii* came with a growth-rate cost in the absence of fluoride, but greater growth rates relative to the ancestors in the presence of fluoride (**Suppl. Fig. 3**).

The fluoride resistance of the bulk populations evolved in the absence of fluoride was also tested in 0.1% glucose (**Suppl. Fig. 6**), and there were no carbon evolution conditions which consistently increased fluoride resistance for *S. mutans* or *S. gordonii*, though there were individual bulk populations that had minor increases in fluoride resistance relative to the ancestor. These observations indicate that the observed fluoride resistance within the populations evolved in the presence of fluoride is likely due to the exposure to fluoride challenge, rather than any collateral effects from adaptation to the alternate carbon environments. Thus, the *S. mutans* populations evolved greater fluoride resistance in fermentable sugars rather than non-fermentable sugar alternatives

### Evolved Isolate & Bulk Population Sugar Alternative Dose Response

The effect of evolving in each sugar alternative on the sensitivity to inhibition by the sugar alternatives (erythritol, sorbitol, xylitol) was tested in the evolved isolates and ancestor strains (**Suppl. Fig. 5**). In *S. mutans*, the area under the curve (AUC) for the evolved population and isolates significantly increased relative to the ancestor for erythritol and xylitol, whereas only the bulk populations increased AUC for sorbitol. These findings indicate that the populations evolved in non-fermentable sugar alternatives (erythritol, xylitol) acquired resistance. Notably, the bulk population evolved in xylitol had a significantly higher AUC than the randomly selected isolated from those populations, again suggesting heterogeneity within the bulk populations. The discrepancy between the *S. mutans* bulk populations and isolates evolved in sorbitol supports the potential heterogeneity in the populations indicated by the fluoride dose response data (**Fig. 2**). Heterogeneity within the *S. mutans* bulk populations increasing resistance to the sugar alternatives could be driven by cooperativity between sub-populations or by a sub-population having higher resistance to sugar alternatives than the randomly selected isolate and overtaking the population when challenged with sugar alternatives in the absence of fluoride. Comparatively, evolved *S. gordonii* bulk populations mildly decreased sensitivity to inhibition by the non-fermentable sugar alternatives (erythritol, xylitol), with notable discrepancies between the bulk populations and isolates in erythritol wherein some isolates did not increase AUC relative to the ancestor. The AUC in sorbitol only increased for one sorbitol bulk population, with mild decreases in AUC in one paired bulk population and isolate. The decreased sorbitol AUC may be due to mutations slowing carbon metabolism, a known mode of acquired fluoride resistance in streptococci^32^.

Taken together, these observations support that there are species specific responses to sugar alternatives and demonstrate the effect of the carbon environment on shaping the available evolutionary trajectories to fluoride resistance.

### Cariogenic Potential of Evolved Isolates

Cariogenic potential of the evolved isolates was assessed using the biofilm forming capabilities and acidogenicity. All isolates evolved in fluoride were tested for their biofilm forming potential in 1% w/v sucrose, glucose, erythritol, sorbitol, or xylitol (**Fig. 3A, Supporting Information**), and their acidogenicity in 0.1% glucose (**Fig. 3B, Supporting Information**). Although there are reports that xylitol resistant clinical isolates are less cariogenic than xylitol sensitive strains^11^, we found that most *S. mutans* isolates evolved in xylitol produced the same amount of biofilm and acid as the ancestor. *S. mutans* strains evolved in sorbitol did not consistently contribute to diminished biofilm formation or any significant shift in acid production, though one of the isolates had decreased biofilm formation in all tested carbon conditions. Interestingly, though some of the isolates evolved in erythritol had moderately diminished biofilm formation in sucrose and glucose, they were the most acidogenic when grown in TSB without dextrose supplemented with 0.1% w/v glucose (adjusted p-value (*p_holm_*)= 5.79 x 10^-3^, 3.68 x 10^-9^, 1.80 x 10^-8^ for Ery,F isolates 1, 2, and 3 respectively). Combined, these observations suggest that *S. mutans* may be driven towards enhanced cariogenicity when evolved in the presence of erythritol.

**Figure 3:**
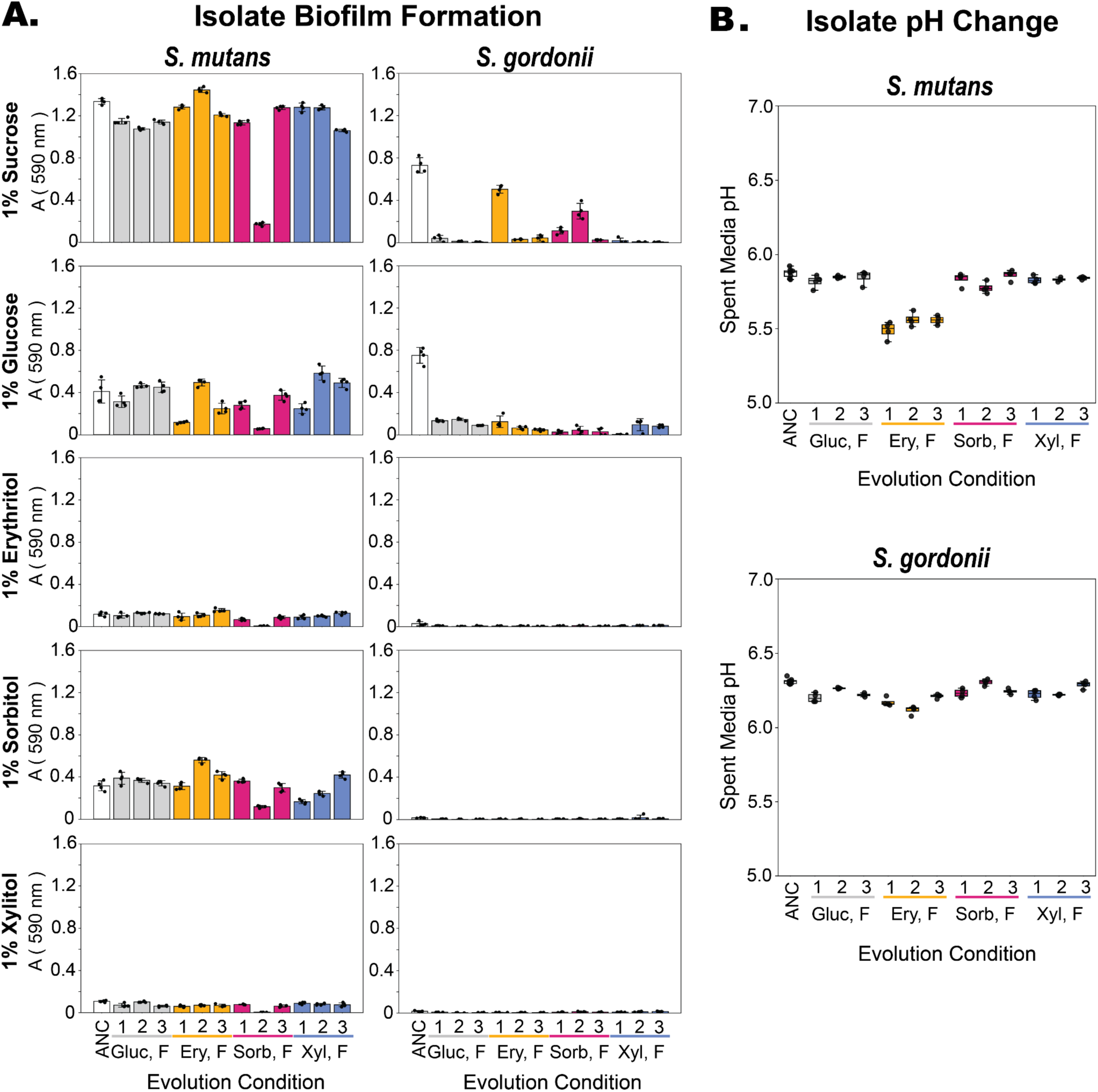
Cariogenic potential of the evolved isolates was investigated using biofilm formation and acid production. (**A**) Biofilm formation over 24 hours of one isolate from each population evolved in fluoride compared to the ancestor strain (ANC) with n=4. Biofilm formation was tested for each strain in 5 different carbon environments (1% w/v sucrose, glucose, erythritol, sorbitol, or xylitol) via crystal violet staining. (**B**) Acidification of base media (TSB without dextrose supplemented with 0.1% w/v glucose) by the ancestors (n=8) and evolved isolates (n=4) was quantified for each isolate using a colorimetric pH indicator. Evolution conditions are as follows: (ANC)= ancestor, (Gluc,F)= 0.1% w/v glucose and sodium fluoride, (Ery, F)= 0.9% w/v erythritol and sodium fluoride, (Sorb, F)= 0.9% sorbitol and sodium fluoride, (Xyl, F)= 0.9% w/v xylitol and sodium fluoride.

Evolved *S. gordonii* strains consistently produced less biofilm than the *S. mutans* strains in glucose and sucrose, and produced little to no detectable biofilm in any of the sugar alternatives. Furthermore, most of the evolved isolates exhibited a drastic reduction in biofilm formation relative to the ancestor, regardless of the evolution carbon environment. There were no drastic shifts in acid production amongst the evolved isolates, though the *S. gordonii* strains evolved in erythritol produced significantly more acid than the ancestor.

### *S. mutans* Isolate Whole Genome Sequencing Analysis

We sent each phenotyped isolate for short-read whole genome sequencing (WGS). Most individuals had multiple mutations in genes previously observed in response to fluoride or sugar alternatives and mutations in novel gene targets.

Xylitol inhibits *S. mutans* growth and biofilm formation by inducing a metabolically costly import-export cycle without being converted into energy in the cell^33^. Many strains with xylitol resistance possess mutations in the phosphotransferase (PTS) system component responsible for xylitol import (smu_872)^34–36^, which stop xylitol import to prevent the costly cycle. Consistent with this ubiquitous mechanism of xylitol resistance, we observed nonsense mutations in smu_872 (fruI) in all *S. mutans* isolates evolved in 0.9% w/v xylitol (**Fig. 4A**).

**Figure 4:**
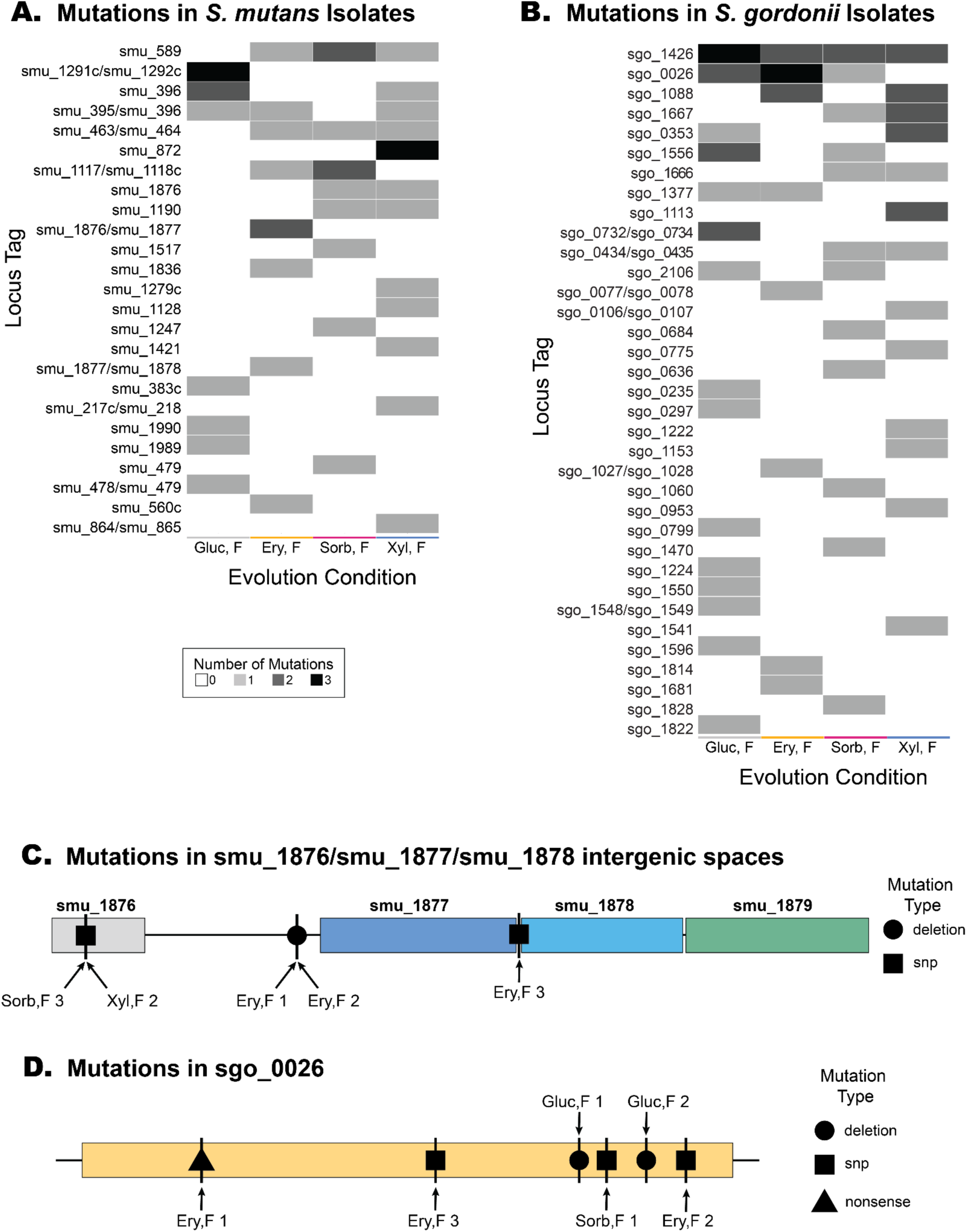
Non-synonymous mutations from the *S.mutans* (**A**) and *S.gordonii* (**B**) evolved isolate whole genome sequencing (WGS) data. Each evolution condition group includes 3 isolates (1 from each replicate population), with evolution conditions as follows: (ANC)= ancestor, (Gluc,F)= 0.1% w/v glucose and sodium fluoride, (Ery F)= 0.9% w/v erythritol and sodium fluoride, (Sorb F)= 0.9% sorbitol and sodium fluoride, (Xyl F)= 0.9% w/v xylitol and sodium fluoride. The heatmap gradient indicates the number of mutations in each locus tag per group. A heatmap including pathway annotations based on the KEGG database for genic regions and maps can be found in **Suppl. Fig. 7**.. Mutation category counts can be found in **Suppl. Tables 7 & 8.** (**C)** Map of the sites that were mutated in *S. mutans* isolates in the smu_1876-smu_1879 region of the genome and (**D**) mutations in *S. gordonii* isolates sgo_0026 (cyclic-di-AMP phosphodiesterase). Mutations are marked by vertical bars, with an overlaid shape indicating the type of mutation at that location (▲=nonsense mutation, ■ = single nucleotide polymorphism, ●= deletion). The isolate(s) that the mutation occurred in is labelled with an arrow pointing to each site.

Clinical interest in erythritol has increased as studies have shown that it can decrease dental biofilm formation^10,12,37^, and erythritol’s mechanisms of inhibition remain under-characterized (though a recent paper proposed a growth inhibition mechanism via targeting celB)^38^. No isolate we evolved in erythritol had a mutation in celB (smu_1596),though some did acquire resistance to erythritol (**Fig. 4A, Suppl. Fig. 5**). An isolate evolved in 0.9% w/v erythritol in the absence of fluoride had a single point mutation (V254I) in a putative ABC Transporter (smu_1934c) but was not mutated in the isolates evolved in erythritol with fluoride. The isolates evolved in erythritol and fluoride had mutations in a histone-like binding protein (smu_589), a DUF1827 family hypothetical protein (smu_560c), and several mutations in intergenic spaces (**Fig. 4A**). Notably, these isolates had mutations in the intergenic spaces surrounding ptnA (smu_1877), which encodes a PTS system EIIAB component responsible for mannose import. Two of the three isolates had the same mutation, a deletion of T at position 1770225 (-106bp downstream of smu_1877 and +771bp upstream of smu_1876, a conserved hypothetical DUF2975 domain-containing protein). The third isolate had a point mutation (A->G, +24/-11) in the intergenic space downstream of smu_1877 and upstream of smu_1878 (ptnC), a mannose-specific PTS system IIC component. The parallel evolution of mutations in these regions, combined with the decreased sensitivity to erythritol in the phenotyping studies, warrants further investigation into the role of ptnA (smu_1877) in erythritol transport and metabolism in *S. mutans* (**Fig. 4C**).

Mutations in the intergenic region upstream of naoX (smu_1117), neighboring an ABC sugar transporter (smu_1118c) occurred in several *S. mutans* isolates evolved in fluoride, including two isolates evolved in sorbitol and one evolved in erythritol. Downregulation of naoX (smu_1117), an NADH oxidase, and smu_1119c have been associated with adaptation to acid shock in *S. mutans*^39,40^.

We observed several mutated genes commonly seen in fluoride resistant *S. mutans* strains^24,32^. For example, a glycerol uptake facilitator protein glpF (smu_396), had mutations (L233F, P235L) in two of the isolates evolved in glucose and fluoride and in one isolate evolved in xylitol and fluoride (E276G). Furthermore, the intergenic space upstream of glpF was mutated in one isolate evolved in glucose and fluoride, one isolate evolved in sorbitol and fluoride, and one isolate evolved in xylitol and fluoride, supporting its potential significance in gene regulation for enhanced fluoride resistance ^24^. Two *S. mutans* isolates evolved in glucose and fluoride had mutations in the intergenic space 38-43bp upstream of chorismate mutase (smu_1291c), a gene that has recently been linked to fluoride resistance in *S. mutans* ^25^. These mutations may be in a regulatory site, potentially increasing the expression level of chorismate mutase and thus increasing fluoride resistance.

Many of the mutated genes lacked KEGG pathway annotations due to limited characterization and/or primarily structural roles. Amongst the annotated genes, the most commonly mutated pathways in the *S. mutans* isolates were glycolysis/gluconeogenesis and the two-component system (**Suppl. Fig. 7**), underscoring the importance of modulating carbon metabolism as a mode of fluoride resistance in *S. mutans*, as previously reported^32^.

### *S. gordonii* Isolate Whole Genome Sequencing Analysis

Enolase (eno, sgo_1426), which converts 2-phosphoglycerate (2-PG) into phosphoenolpyruvate (PEP) during glycolysis, was the most commonly mutated gene across *S. gordonii* isolates (**Fig. 4B, Suppl. Fig. 7 & 8, Suppl. Table 6**). This observation aligns with previous studies which observed enolase mutations in fluoride resistant streptococci^24,32^ and studies demonstrating it is commonly upregulated by fluoride riboswitches^41^. Notably, while previous studies identified mutations along the octamer interface of the enolase^42^ in fluoride resistant *S. mutans*, the most common mutation in our *S. gordonii* dataset (T289I) was near the metal cofactor binding in the active site. The mutation T289I in sgo_1426 was present in at least one isolate evolved in each carbon environment, indicating its fluoride-specific effect likely protecting from fluoride challenge by perturbing the active site. The homologous mutation, T287I in *S. mutans* enolase (smu_1247), was initially observed in an *in vitro* evolution experiment with fluoride^24^ and was present in one *S. mutans* isolate evolved in sorbitol and fluoride in this experiment. These enolase mutations are predicted to decrease enolase inhibition by fluoride.

The second most commonly mutated gene in *S. gordonii* was sgo_0026, which encodes a cyclic-di-AMP phosphodiesterase responsible for regulating osmotic stress, acid tolerance, and cell wall homeostasis^43^. Though sgo_0026 was mutated in 5 isolates, the mutation type and site were inconsistent, including substitutions at various sites, frameshift, and a non-sense mutation (**Fig. 4D**). The mutations in sgo_0026 likely result in gene inactivation and/or lowered catalytic efficiency based on their positions in the DDH and DHHA1 domains.

In environments with xylitol and fluoride, sgo_113-the homologue of *S. mutans’* xylitol importer (smu_872)-was mutated in 2 out of the 3 *S. gordonii* isolates. Notably, these mutations were non-synonymous point mutations (Q190P, G299D) rather than the nonsense mutations in the *S. mutans* isolates.

At the pathway level, teichoic acid biosynthesis (sgo_1377, sgo_0775), purine metabolism, and carbon metabolism (glycolysis/gluconeogenesis, starch and sucrose metabolism) were most perturbed across *S. gordonii* isolates (**Suppl. Fig. 7**). Thus, similarly to *S. mutans,* slowing central carbon metabolism is likely one of the primary protective mechanisms to fluoride challenge for *S. gordonii*.

### Mutated histone-like binding protein increases fluoride resistance in *S. mutans*

A histone-like binding protein (HLP, smu_589) was the most frequently mutated gene in the evolved *S. mutans* isolates. Notably, HLP is the only nucleoid-associated protein identified in *S. mutans* and is an essential gene in streptococci ^46^. Streptococcal HLPs, homologous to Huβ, have an essential role in global DNA architecture, gene transcription, and can have immunogenic effects when secreted^47–49^. Four *S. mutans* isolates evolved in fluoride and a sugar alternative (0.9% w/v erythritol, sorbitol, or xylitol) had mutations in this gene. Neither smu_589, nor its homologue (sgo_0701), were mutated in *S. mutans* isolates evolved in fluoride with glucose or in *S. gordonii* isolates evolved in any condition

The most common mutation in smu_589, K4Q, occurred in 3 isolates, one from each of the sugar alternative treatments in the presence of fluoride. One other mutation in a DNA-binding region, R62S, occurred in an isolate evolved in 0.9% w/v sorbitol and fluoride. HLP proteins generally function as a homodimer, with the lysine residues responsible for non-specific DNA binding and dimerization. Lysine 4 in *S. mutans*’ HLP protein aligns with Lysine 3 in the most commonly studied Huβ, and is one of the most highly conserved residues across species, impacting both protein dimerization and DNA-binding in homologous strains^49,50^. Lysine acetylation serves as a regulatory mechanism to modulate HLP-DNA binding and overall DNA compression to modulate the accessibility for transcription. The observed mutation, K4Q, mimics lysine acetylation by removing the positive charge, likely impacting the dimerization of the protein, DNA-binding, and thus altering gene expression. To our knowledge, this mutation has not been previously reported in clinically derived fluoride resistant strains.

To characterize the effect of this mutation on fluoride sensitivity, the *S. mutans* ancestor background was transformed with the K4Q point mutation in smu_589. The K4Q mutant had significantly increased fluoride resistance, with minor effects on growth in the presence of sugar alternatives (**Fig. 5**). Notably, the K4Q mutation significantly shifted fluoride sensitivity in 0.1% w/v glucose with the IC_50_ increasing from 4.2 ± 0.2 mM to 4.9 ± 0.2 mM (p-value < 0.001), though the largest increase fluoride resistance occurred in 0.9% w/v Xylitol the IC_50_ increasing from 4.0 ± 0.4 mM to 6.4 ± 0.3 mM (p-value < 0.001). This trend held in 0.9% w/v erythritol but we did not observe a significant difference from the ancestor in 0.9% w/v sorbitol (**Fig. 5B, Suppl. Fig. 7).** When growth curves were compared to the WT ancestor, the K4Q mutation showed a cost in the absence of fluoride in, as shown by decreased maximum growth rate (μ_max_) in 0.1% w/v gluose and 0.9% w/v sorbitol and increased lag time in all carbon conditions (**Fig. 5C & D**). However, when strains were grown in 2 mM fluoride, the K4Q mutation was beneficial, as shown by increased μ_max_ relative to the WT ancestor in 0.9% w/v erythritol and 0.9% xylitol and decreased lag time relative to the ancestor in all carbon conditions **(Fig. 5C& D, Suppl. Fig. 9**). These observations could be due to K4Q reducing HLP-mediated DNA compaction, leading to changes in gene expression similar to a broad stress response which aid increased fluoride resistance.

**Figure 5:**
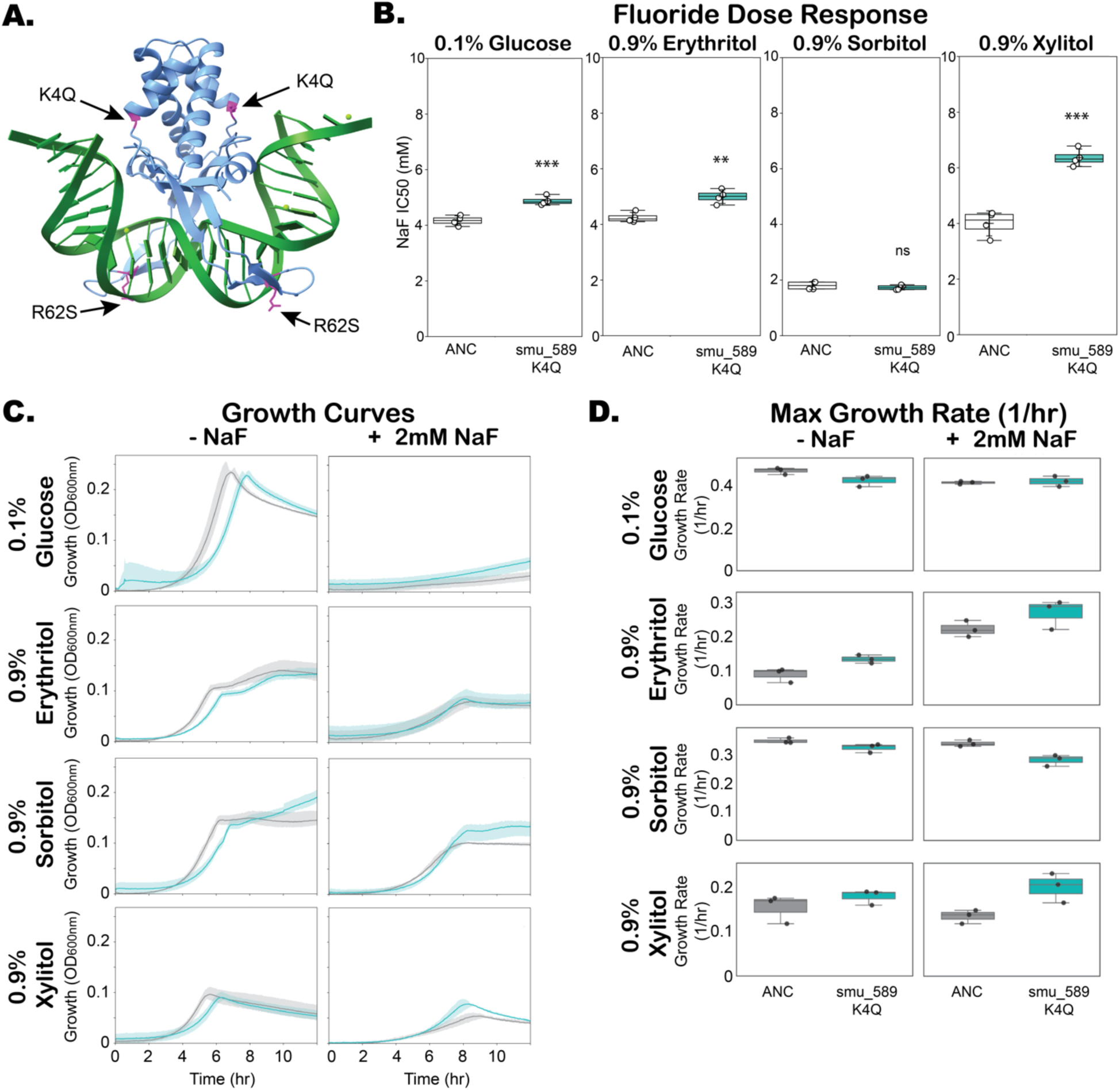
Parallel evolution of mutations in a histone-like binding protein (HLP) in *S. mutans* isolates resulted in increased fluoride resistance. (**A**) Alphafold3 model of an smu_589 (HLP) homodimer (blue) bound to a DNA fragment (green), with the locations of the mutated residues highlighted in magenta and annotated with arrows ^44,45^. (**B**) Fluoride IC_50_ values for the ancestor (ANC) and the ancestor strain with K4Q point mutation in smu_589 (smu_589 K4Q). Values were fit from dose response curves (n=3). Fluoride mean IC_50_ ± SD is reported in **Suppl. Table 9**. For significance: ns= not significant, *= p-value <0.05, **= p-value <0.01, ***= p-value <0.001. (**C**) Growth of the ancestor (grey) and ancestor with the smu_589 K4Q mutation (teal) over 12 hours (n=3), with the mean line surrounded by the range (shaded region). Each strain was grown in the absence (- NaF) and presence (+ 2mM NaF) of fluoride in 4 carbon environments. 24-hour growth curve data compared to evolved isolate growth curved can be found in **Suppl. Fig. 9**. (**D)** Maximum growth rate of the ancestor and smu_589 mutant over the 24-hour growth curve (n=3). Mean ± SD for the time to maximum growth rate and maximum growth rates can be found in **Suppl. Materials**.

## Conclusions

We investigated the impact of carbon environment on fluoride resistance via the *in vitro* evolution of *S. mutans* and *S. gordonii* in commonly used sugar alternatives (**Fig. 1**). The presence of fermentable versus non-fermentable sugars impacted the efficiency and mode of fluoride resistance in cariogenic *S. mutans*, but had little effect on the degree of fluoride resistance in the commensal *S. gordonii* (**Fig. 2**). These findings are particularly relevant in light of the growing interest in the use of non-fermentable sugar alternatives in combination with fluoride to manage caries, as non-fermentable sugar alternatives may decrease the adaptability of a core cariogenic strain in ECC. When biofilm formation potential and acid production were assessed as proxies for cariogenic potential, evolving in the presence of non-fermentable sugar alternatives had minimal effects on biofilm formation potential of *S. mutans* strains. However, *S. mutans* acidogenesis significantly increased when evolved in erythritol, indicating that erythritol may have less utility as a prebiotic than xylitol for clinical applications. Notably, evolving in the presence of the most commonly used oral antimicrobial agent, fluoride, decreased the biofilm formation potential of *S. gordonii* strains relative to the ancestor regardless of carbon condition (**Fig. 3**).

Mechanisms of acquired fluoride resistance varied both between *S. mutans* and *S. gordonii* and within the same species, as few genes were mutated in multiple isolates. Our findings replicated some of the most commonly mutated genes conferring fluoride resistance in clinical isolates, such as a glycerol uptake facilitator (glpF, smu_396) and enolase (eno, smu_1247/sgo_1426), largely associated with decreased metabolism. Additionally, we observed multiple novel mutations conferring fluoride resistance, such as in a key mutation in one of the most conserved sites (Lysine 4) in the only reported nucleoid associated protein in streptococci, HLP (smu_589). We also identified smu_1877 (ptnA) as a PTS component potentially involved in erythritol import in *S. mutans* due to multiple mutations in a small upstream intergenic region which may act as a regulatory site for smu_1877 expression. Together, these findings underscore the importance of understanding the impact of carbon source on the evolutionary trajectories of cariogenic and non-cariogenic microbiota and may inform the optimization of the use of sugar alternatives in combination with anti-caries therapies.

We have established that non-fermentable sugar alternatives can limit the degree of acquired fluoride resistance in *S. mutans,* though chronic exposure and evolutionary adaptation to some, such as erythritol, may also drive enhanced acid production and thus cariogenic potential (**Fig. 3B**). Further investigation into the long-term effect of other sugar alternatives (e.g., D-tagatose, allulose, aspartame) as potential adjuvants for caries prevention and treatment is still needed. We also have identified several previously unreported mutations which may contribute to fluoride resistance, which could be examined in clinical sequencing datasets (**Fig. 4AB**). Notably, the mechanism by which the most common mutation (K4Q) in *S. mutans*’ histone-like binding protein increases *S. mutans* fluoride resistance warrants further investigation. Finally, we acknowledge that *in vivo* evolution does not occur in isolation; instead, evolution occurs in diverse communities shaped by numerous interspecies interactions, many of which are driven by nutrient availability. The effects of sugar alternatives on the cariogenic strains, such as *S. mutans*, may vary greatly in these communities, and may also impact their evolutionary trajectories in response to common environmental stressors such as fluoride.

## Author Contributions

M.R.S. Conceptualization, assay design, data collection, data processing & analysis, writing-original draft

M.A.E. Data collection, writing-review & editing

R.B.S. Funding acquisition, writing-review & editing

L.M.A.T. Funding acquisition, writing-review & editing

X.N.L. Conceptualization, writing-review & editing

K.B.W. Funding acquisition, conceptualization

L.Z. Funding acquisition, conceptualization, writing-review & editing

## Acknowledgements

This work was supported by NIH T32 GM145470 (to M.R.S.) and NIH RM1 DE034220 (to R.B.S., L.M.A.T., K.B.W., L.Z.). We thank members of the Wood, Zaman, Lin, Tenuta, and Stockbridge laboratories for helpful discussions.

This work would not have been possible without Dr. Kevin B. Wood’s thoughtful mentorship and infectious excitement about research, we are very grateful for his continued impact.

## Data Archiving Statement

Data for this study are available at: *to be completed after manuscript is accepted for publication*. All data used in this work will be archived and publicly available at [Deep Blue Data](https://deepblue.lib.umich.edu/data) upon publication. These data will include the raw data for dose response curves, growth curves, pH determination, and biofilm formation as well as tables of the fit values and summary statistics for Inhibitory Concentration 50% (IC_50_) values. Code used to analyze these data and generate figures will be available on GitHub.

## Notes

### Competing Interest Statement

The authors have declared no competing interest.

