## Supplementary material for "Sugar alternatives shift the evolutionary trajectory of fluoride resistance in cariogenic and non-cariogenic dental bacteria": [Read Me].rtf

Supplemental Materials
This document contains additional methods, figures, and tables referenced directly in the main text. 

Data Availability:
All data,  code used for analyses, and strains will be available upon request to the authors. 

Please note that raw data will be archived on Deep Blue Data. [links will be added prior to publication]

Processed data & statistics are available here in the “Supplemental Data & Statistics” folder. If you have any difficulty finding what you are looking for please contact the authors. [contact information added prior to publication]

P.S. If you are reading this— thanks for reading the supplemental, I hope that you enjoyed the paper. 
