## Supplemental Materials for "Sugar alternatives shift the evolutionary trajectory of fluoride resistance in cariogenic and non-cariogenic dental bacteria"

### Supplemental Methods ........................................................... 2-3

Supplemental Method 1: Natural Transformation in *S.mutans* UA159 ................................ 2-3

### Supplemental Data ........................................................... 3-16

**Supplemental Method 1:** Natural Transformation in *S. mutans* UA159

Day 0:

1. Prepare fresh streaks of *S. mutans* on TSB with dextrose agar plates
    a. Place at 37°C w/ 5% $CO_2$ overnight (at least 16 hours) to get juicy colonies
2. Prepare TSB with dextrose agar plates w/ any selective agent you may be using
3. Prepare 30% glycerol stock
4. Prepare CDM
    a. Note: CDM should be prepared fresh weekly, and discarded earlier if there is any visible precipitation
5. Prepare XIP solution (1mM XIP)
    a. XIP: NH2-GLDWWSL-COOH
        i. Note: we ordered from GenScript, but you can order from any supplier)
    b. Prepare solution in MQ water and store in -20°C

Day 1:

6. Condition cells by inoculating 3mL of CDM w/ 1% glucose with fresh colonies
    a. Grow static at 37C w/ 5% $CO_2$
    b. Periodically monitor $OD_{600nm}$
    c. Once cells reach $OD_{600nm}$ 0.5 go to step 2
        i. Note: likely will take several hours, growth is slow in CDM
7. Prepare conditioned cell stock
    a. Dilute fresh culture 1:1 w/ 30% v/v glycerol (final glycerol 15% v/v)
    b. Flash freeze cells & store in –80°C

Day 2:

8. Revive conditioned cells
    a. Dilute conditioned cells (from –80°C) in CDM 1:10
        i. I do ~2 mL of culture, but this volume is flexible
        ii. Also make a blank for OD measurements
    b. Incubate at 37°C w/ 5% $CO_2$ static until cells reach $OD_{600nm}$=0.1
9. Induce transformation
    a. Aliquot culture into sterile 1.5 mL tubes
        i. I do ~0.5-1 mL typically
    b. Add DNA (final concentration 0.1-1 ug/mL) and XIP (1 uM XIP)
        i. Note: for the stock concentration of XIP I work with (1 mM XIP) add 1 uL
        ii. Note: DNA concentration will vary for plasmid, the value shown above is for linear fragments
    c. Incubate for ~3 hr at 37°C in closed tubes
        i. Note: do not incubate less than 2 hr, efficiency declines significantly
10. Plate on TSB dextrose agar with any relevant selector (ie antibiotic)
    a. Grow at 37°C w/ 5% $CO_2$ for 20-24 hr
11. Screen for mutation with colony PCR

    a. When setting up the colony PCR, spot 2 uL of the water/cell slurry into a fresh TSB dextrose plate (with any appropriate selector)

12. Re-streak colonies with PCR-confirmed mutation onto TSB dextrose plate

a. <u>Note:</u> if doing markerless mutagenesis (ie no selective plating), we recommend repeating steps 11-12 multiple times to ensure you have a pure mutant.

13. Confirm mutations w/ Sanger Sequencing

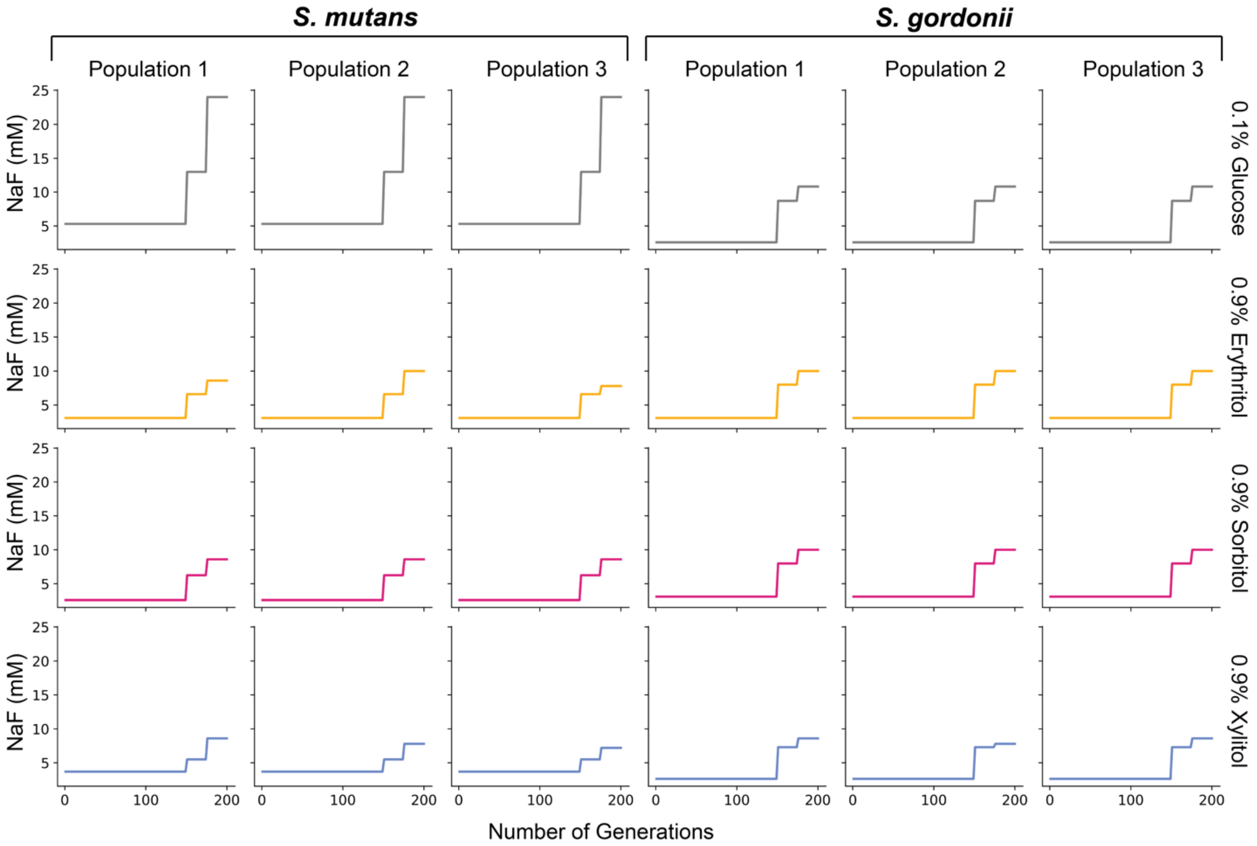

**Supplemental Figure 1:** Sodium Fluoride concentration (mM) over the course of the *in vitro* evolution for each population.

**Supplemental Table 1:** Primers used to screen for the K4Q mutation in *S. mutans* colonies after transformation & to send region for Sanger sequencing to confirm the mutation.

| **Application** | **Target** | **FWD Primer** | **REV Primer** |
|---|---|---|---|
| Homologous Recombination | 3.3 kbp region including smu_589 | GCTGGAGCAGATGTCACTGT | AAAAGGGGAGTCTCGCAGC |
| Colony PCR Screening | smu_589 WT | TGGAGGATTTGTTAAATGGCT AACA | TTTGAGGGTTGCGACCTTTG |
| Colony PCR Screening | smu_589 K4Q | TGGAGGATTTGTTAAATGGCT AAGC | TTTGAGGGTTGCGACCTTTG |
| Sanger Sequencing | smu_589 region | CTGCCAGCATCAGAGGTTCT | TTTGAGGGTTGCGACCTTTG |

**Supplemental Table 2:** Summary statistics for the one-way ANOVA and Tukey HSD comparison of the fluoride $IC_{50}$ of the *S. mutans* bulk populations and isolates evolved in fermentable (glucose and sorbitol) and non-fermentable (erythritol and xylitol) in the presence of fluoride. One-way ANOVA: $F(4,84) = 33.2$, $p=1.36 \times 10^{-22}$. For significance: ns= not significant, *= adjusted p-value <0.05, **= adjusted p-value <0.01, ***= adjusted p-value <0.001.

| Group 1 | Group 2 | Mean difference (mM) | Adjusted p-value | CI95 lower limit | CI95 upper limit | Reject null ? | significance |
|---|---|---|---|---|---|---|---|
| Fermentable-bulk | *S. mutans* ancestor | -17.58 | 4.52E-12 | -23.30 | -11.85 | TRUE | *** |
| Fermentable-isolate | *S. mutans* ancestor | -11.31 | 1.35E-06 | -16.78 | -5.83 | TRUE | *** |
| Non-fermentable-bulk | *S. mutans* ancestor | -4.01 | 2.88E-01 | -9.67 | 1.65 | FALSE | ns |
| Non-fermentable-isolate | *S. mutans* ancestor | -1.88 | 8.74E-01 | -7.35 | 3.60 | FALSE | ns |
| Fermentable-bulk | Fermentable-isolate | -6.27 | 2.74E-03 | -10.91 | -1.63 | TRUE | ** |
| Fermentable-bulk | Non-fermentable-bulk | -13.57 | 1.60E-10 | -18.42 | -8.71 | TRUE | *** |
| Fermentable-bulk | Non-fermentable-isolate | -15.70 | 1.15E-13 | -20.34 | -11.06 | TRUE | *** |
| Fermentable-isolate | Non-fermentable-bulk | -7.30 | 2.41E-04 | -11.86 | -2.73 | TRUE | *** |
| Fermentable-isolate | Non-fermentable-isolate | -9.43 | 3.49E-07 | -13.76 | -5.10 | TRUE | *** |
| Non-fermentable-bulk | Non-fermentable-isolate | -2.13 | 6.90E-01 | -6.70 | 2.43 | FALSE | ns |

**Supplemental Table 3:** Summary statistics for the one-way ANOVA and Tukey HSD comparison of the fluoride $IC_{50}$ of the *S. gordonii* bulk populations and isolates evolved in fermentable (glucose and sorbitol) and non-fermentable (erythritol and xylitol) in the presence of fluoride. One-way ANOVA: $F(4, 80)=55.6$, $p=2.38 \times 10^{-22}$. For significance: ns= not significant, *= adjusted p-value <0.05, **= adjusted p-value <0.01, ***= adjusted p-value <0.001.

| Group 1 | Group 2 | Mean difference | Adjusted p-value | CI95 lower limit | CI95 upper limit | Reject null ? | significance |
|---|---|---|---|---|---|---|---|
| Fermentable-bulk | *S. gordonii* ancestor | -9.26 | 1.61E-14 | -11.20 | -7.32 | TRUE | *** |
| Fermentable-isolate | *S. gordonii* ancestor | -8.16 | 1.61E-14 | -10.05 | -6.26 | TRUE | *** |
| Non-fermentable-bulk | *S. gordonii* ancestor | -8.17 | 1.61E-14 | -10.10 | -6.24 | TRUE | *** |

| | | | | | | | |
|---|---|---|---|---|---|---|---|
| Non-fermentable-isolate | *S. gordonii* ancestor | -6.01 | 1.31E-12 | -7.90 | -4.13 | TRUE | *** |
| Fermentable-bulk | Fermentable-isolate | -1.10 | 5.92E-02 | -2.22 | 0.03 | FALSE | ns |
| Fermentable-bulk | Non-fermentable-bulk | -1.09 | 8.46E-02 | -2.27 | 0.09 | FALSE | ns |
| Fermentable-bulk | Non-fermentable-isolate | -3.24 | 3.36E-11 | -4.35 | -2.14 | TRUE | *** |
| Fermentable-isolate | Non-fermentable-bulk | 0.01 | 1.00E+00 | -1.10 | 1.12 | FALSE | ns |
| Fermentable-isolate | Non-fermentable-isolate | -2.15 | 1.15E-06 | -3.17 | -1.12 | TRUE | *** |
| Non-fermentable-bulk | Non-fermentable-isolate | -2.16 | 3.74E-06 | -3.24 | -1.07 | TRUE | *** |

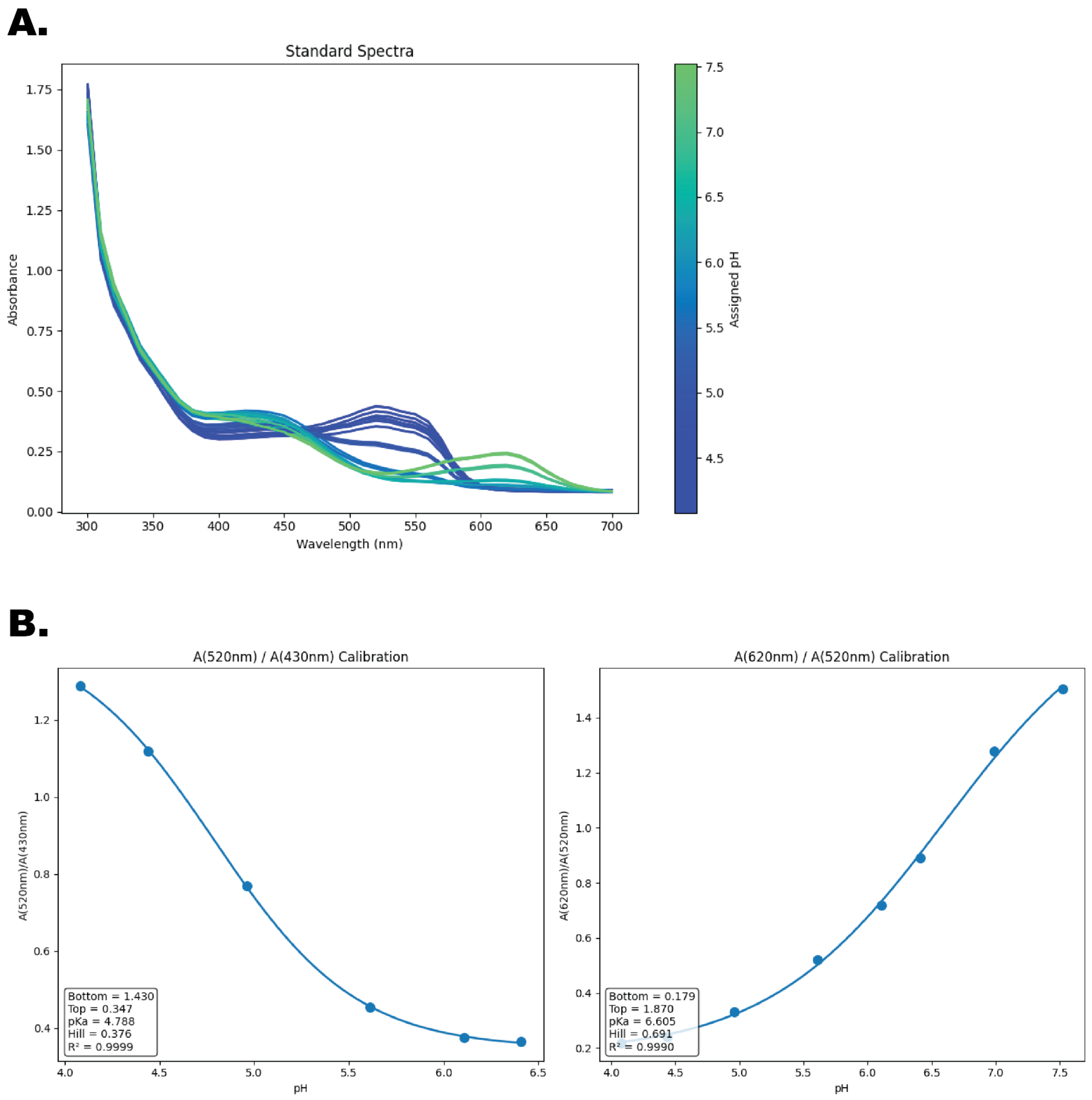

**Supplemental Figure 2:** (**A**) Absorbance spectra of pH calibrated TSB without dextrose supplemented with 0.1% w/v glucose after addition of the colorimetric pH indicator dye. Color of each curve is based on the pH of the control, with an n=4. (**B**) The absorbance ratios of the top peaks with pH shifts fit to a 4-parameter logistic curve. Note that the curve was fit to the mean of the 4 replicates.

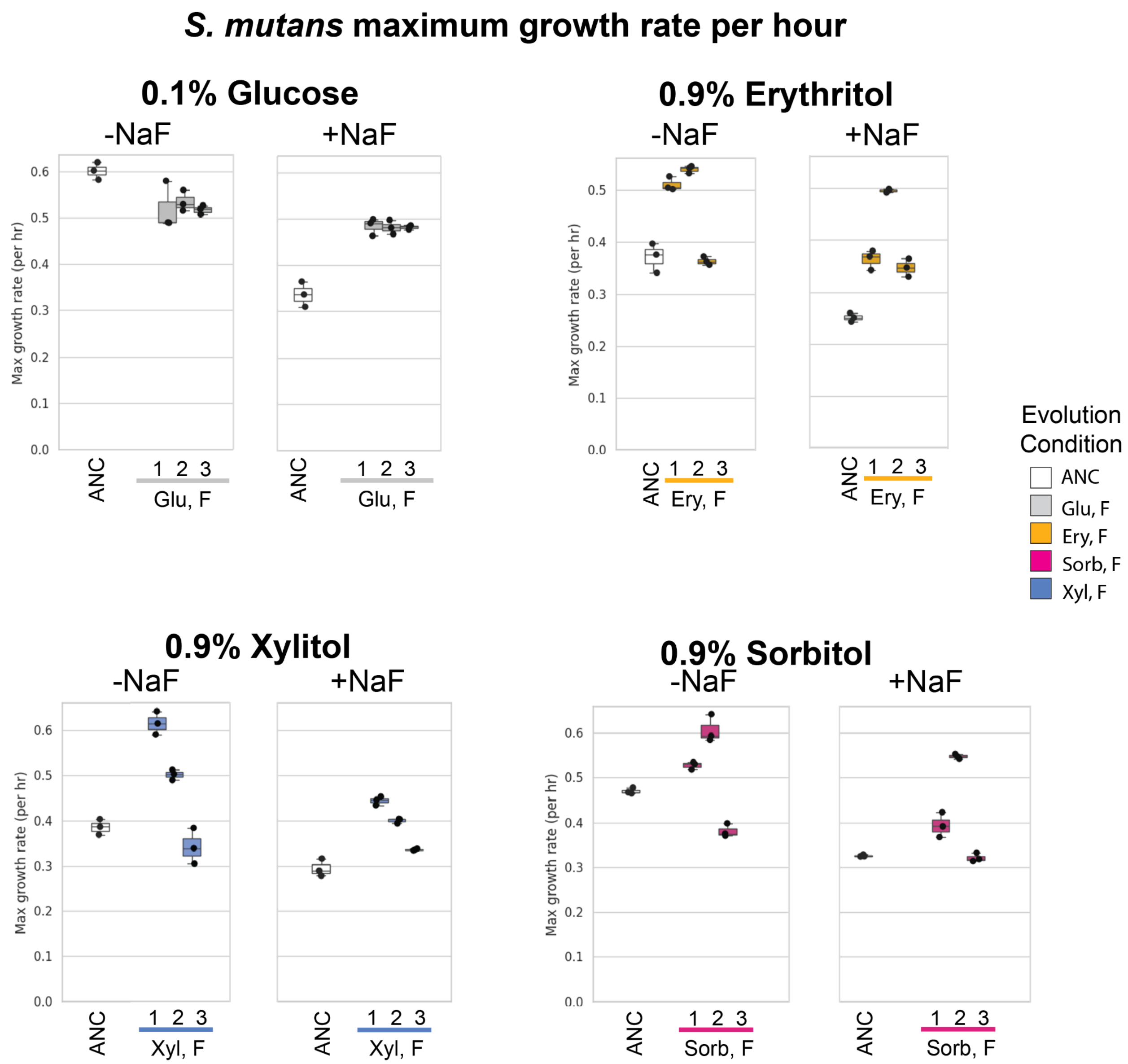

**Supplemental Figure 3:** Maximum growth rate ($\mu_{max}$) per hour of the evolved isolates compared to the ancestor (ANC) in the carbon environment they evolved in. The maximum growth rate was determined when grown in the absence of fluoride and in the presence of 2mM NaF with n=3. Evolution conditions are as follows: (ANC)= ancestor, (Gluc,F)= 0.1% w/v glucose and sodium fluoride, (Ery, F)= 0.9% w/v erythritol and sodium fluoride, (Sorb, F)= 0.9% w/v sorbitol and sodium fluoride, (Xyl, F)= 0.9% w/v xylitol and sodium fluoride.

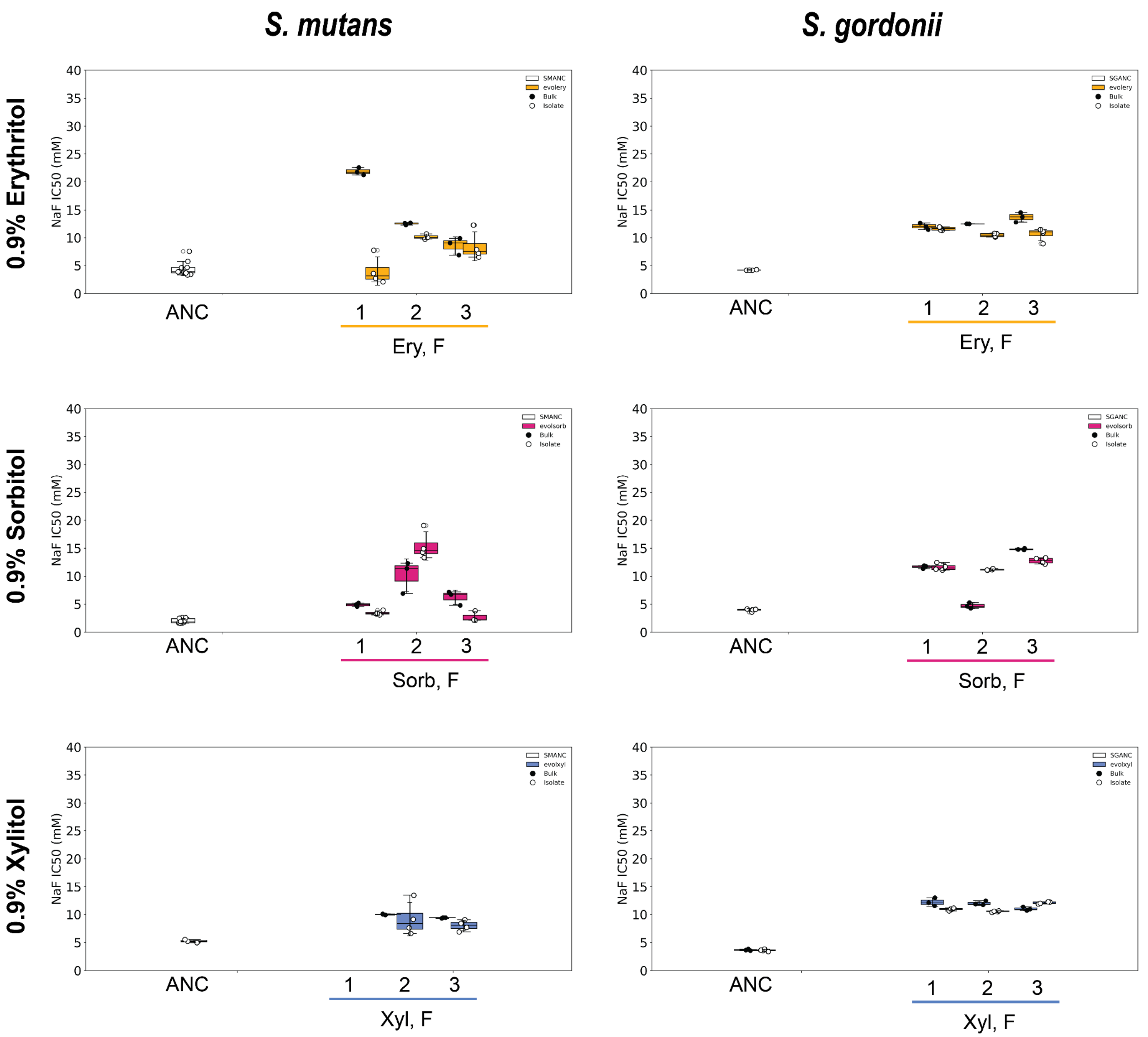

**Supplemental Figure 4:** Fluoride $IC_{50}$ of the 3 replicate bulk populations and randomly selected isolates evolved in sugar alternatives for *S. mutans* (left) and *S. gordonii* (right) when tested in the carbon environment they were evolved in with n=3 for the bulk and n=4 for the isolates. Evolution conditions are as follows: (ANC)= ancestor, (Ery F)= 0.9% w/v erythritol and sodium fluoride, (Sorb F)= 0.9% sorbitol and sodium fluoride, (Xyl F)= 0.9% w/v xylitol and sodium fluoride.

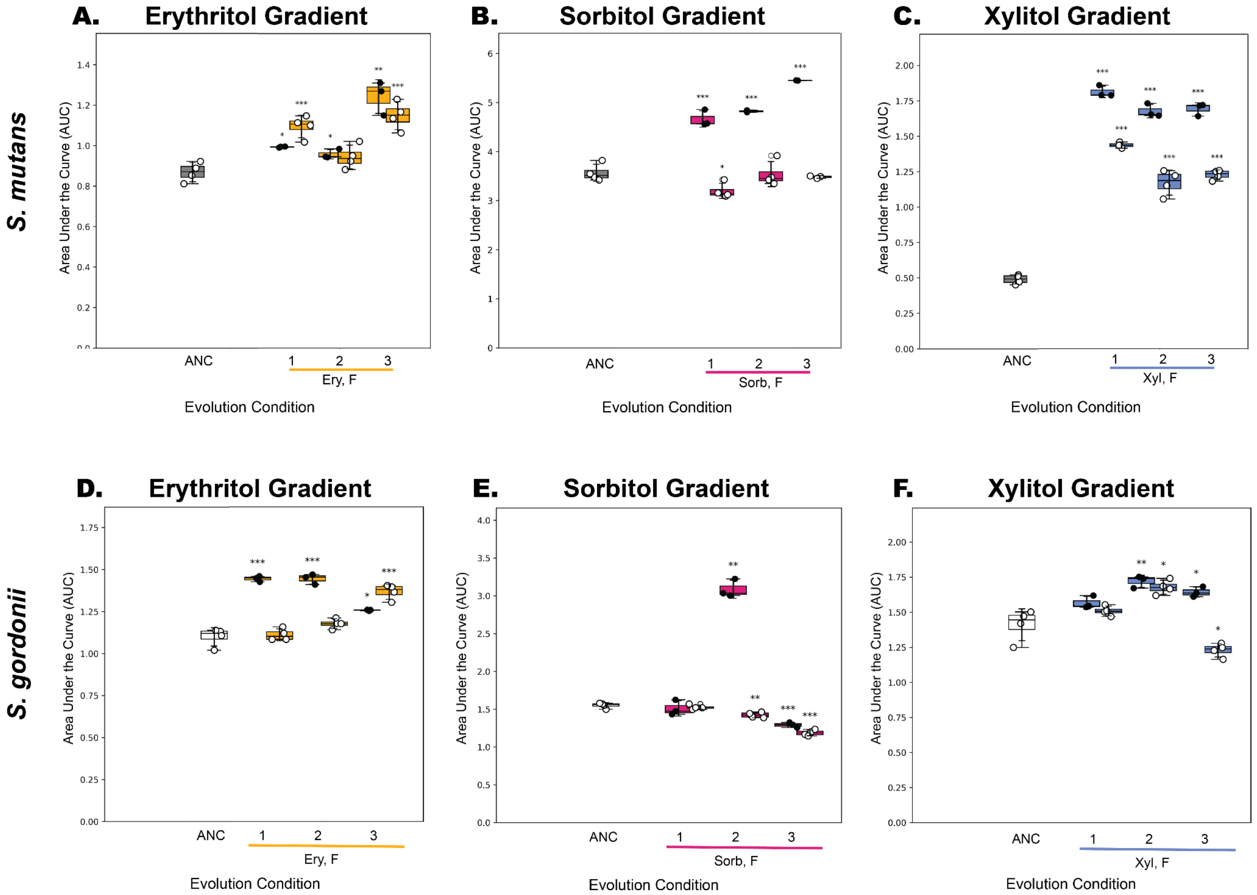

**Supplemental Figure 5:** Area under the curve (AUC) for the growth of the bulk populations (●, n=3) and isolates (○, n=4) grown in a gradient of sugar alternatives (2 fold serial dilution from 10% w/v to 0% w/v). The sugar alternative that was tested is labelled at the top of each plot, with the evolution condition of the bulk and isolate populations labelled on the x axis as follows: (ANC)= ancestor, (Gluc, F)= 0.1% w/v glucose and sodium fluoride, (Ery, F)= 0.9% w/v erythritol and sodium fluoride, (Sorb, F)= 0.9% w/v sorbitol and sodium fluoride, (Xyl, F)= 0.9% w/v xylitol and sodium fluoride. The top row (A, B,C) are *S. mutans*, whereas the bottom row (D,E,F) are *S. gordonii.* For significance: ns= not significant, *= adjusted p-value <0.05, **= adjusted p-value <0.01, ***= adjusted p-value <0.001.

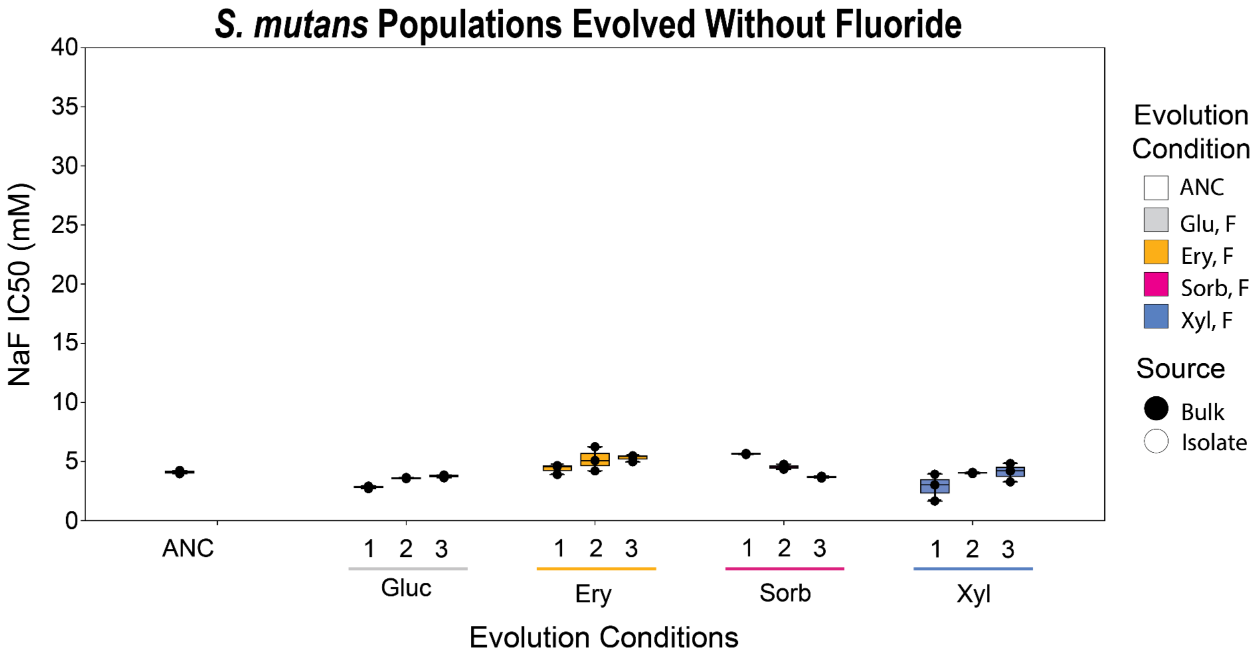

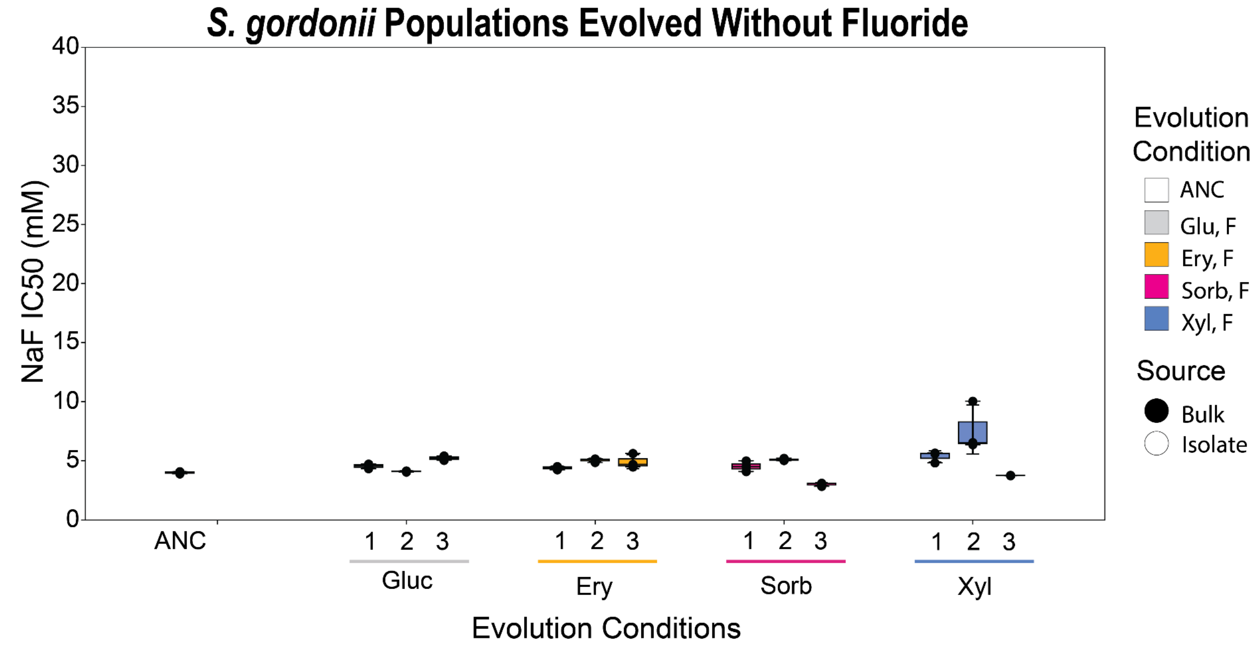

**Supplemental Figure 6:** Fluoride $IC_{50}$ for the 3 replicate bulk populations in sugar alternatives from *S. mutans* (top) and *S. gordonii* (bottom) when tested in 0.1% glucose with n=3. Evolution conditions are as follows: (ANC)= ancestor, (Gluc)= 0.1% w/v glucose, (Ery)= 0.9% w/v erythritol, (Sorb)= 0.9% w/v sorbitol, (Xyl)= 0.9% w/v xylitol.

**Supplemental Table 4:** Fluoride $IC_{50}$ for the 3 replicate bulk populations in sugar alternatives from *S. mutans* when tested in 0.1% glucose with n=3. Evolution conditions are as follows: (ANC)= ancestor, (Gluc)= 0.1% w/v glucose, (Ery)= 0.9% w/v erythritol, (Sorb)= 0.9% w/v sorbitol, (Xyl)= 0.9% w/v xylitol.

| **Population Number** | **Evolution Condition** | **$IC_{50}$ ± SD (mM)** |
|---|---|---|

| | | |
|---|---|---|
| | None (*S.mutans* ANC) | 4.10 ± 0.14 |
| 1 | Gluc | 2.84 ± 0.12 |
| 2 | Gluc | 3.59 ± 0.05 |
| 3 | Gluc | 3.75 ± 0.12 |
| 1 | Ery | 4.38 ± 0.41 |
| 2 | Ery | 5.17 ± 1.03 |
| 3 | Ery | 5.29 ± 0.28 |
| 1 | Sorb | 5.64 ± 0.05 |
| 2 | Sorb | 4.53 ± 0.21 |
| 3 | Sorb | 3.70 ± 0.09 |
| 1 | Xyl | 2.86 ± 1.14 |
| 2 | Xyl | 4.04 ± 0.06 |
| 3 | Xyl | 4.10 ± 0.78 |

**Supplemental Table 5:** Fluoride $IC_{50}$ for the 3 replicate bulk populations in sugar alternatives from *S. gordonii* when tested in 0.1% glucose with n=3. Evolution conditions are as follows: (ANC)= ancestor, (Gluc)= 0.1% w/v glucose, (Ery)= 0.9% w/v erythritol, (Sorb)= 0.9% w/v sorbitol, (Xyl)= 0.9% w/v xylitol.

| Population Number | Evolution Condition | IC50 ± SD |
|---|---|---|
| | None (*S.gordonii* ANC) | 3.99 ± 0.08 |
| 1 | Gluc | 4.54 ± 0.19 |
| 2 | Gluc | 4.10 ± 0.05 |
| 3 | Gluc | 5.22 ± 0.19 |
| 1 | Ery | 4.39 ± 0.14 |
| 2 | Ery | 5.04 ± 0.16 |
| 3 | Ery | 4.90 ± 0.61 |
| 1 | Sorb | 4.52 ± 0.45 |
| 2 | Sorb | 5.09 ± 0.10 |
| 3 | Sorb | 3.02 ± 0.15 |
| 1 | Xyl | 5.35 ± 0.47 |

| | | |
|---|---|---|
| 2 | Xyl | 7.64 ± 2.07 |
| 3 | Xyl | 3.75 ± 0.01 |

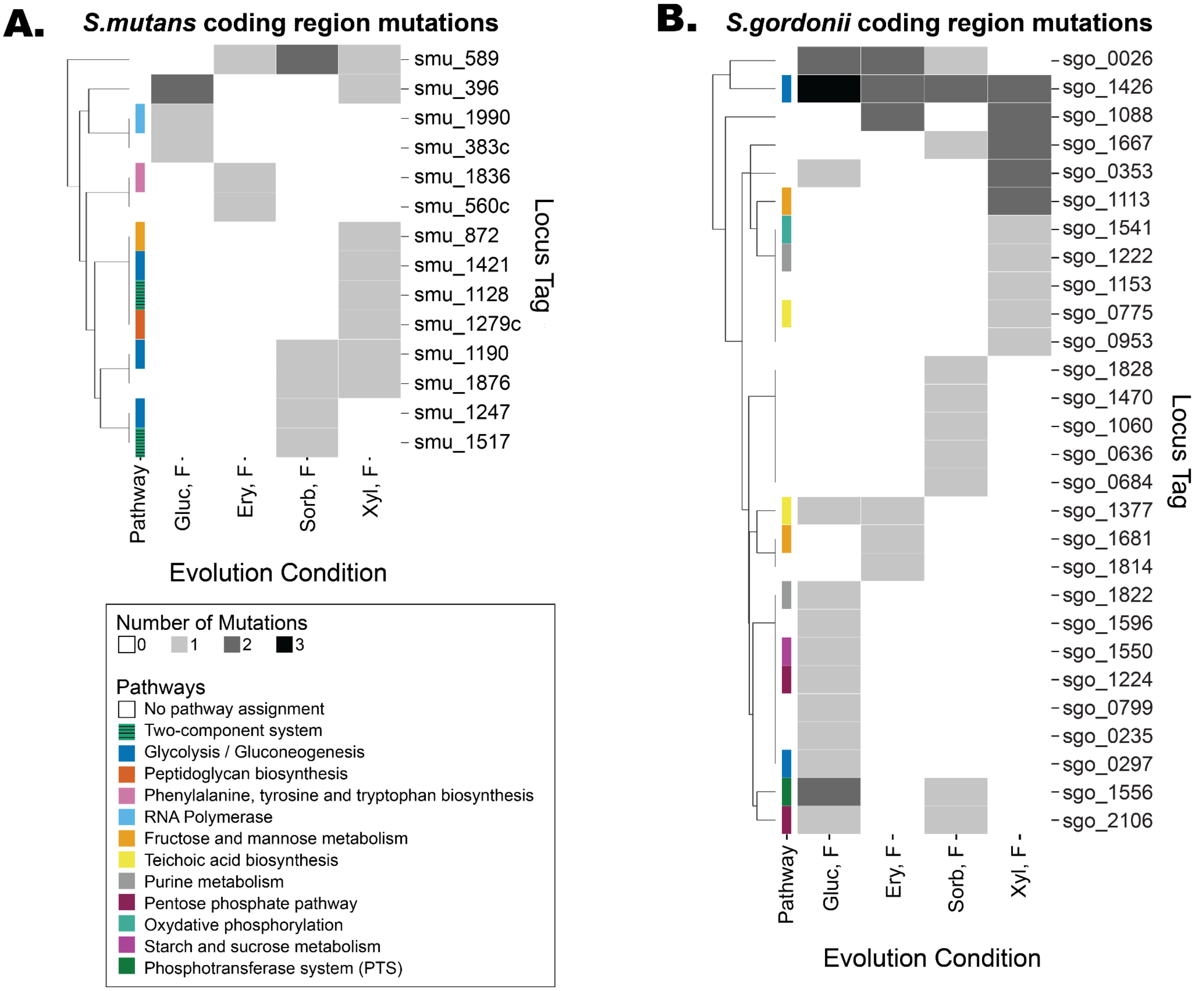

**Supplemental Figure 7:** Heatmaps of the mutated coding regions from the *S.mutans* (**A**) and *S.gordonii* (**B**) isolate whole genome sequencing (WGS) data. Each evolution condition group includes 3 isolates (1 from each replicate population), with the gradient indicating the number of mutations in each locus tag per group. Only locus tags for genic regions are included in this figure, an expanded heatmap including intergenic region mutations is **Figure 4**. Locus tags are clustered based on how commonly genes were mutated simultaneously. The Pathway bar on the heatmap indicates the pathway assignment, when available, based on the KEGG database.

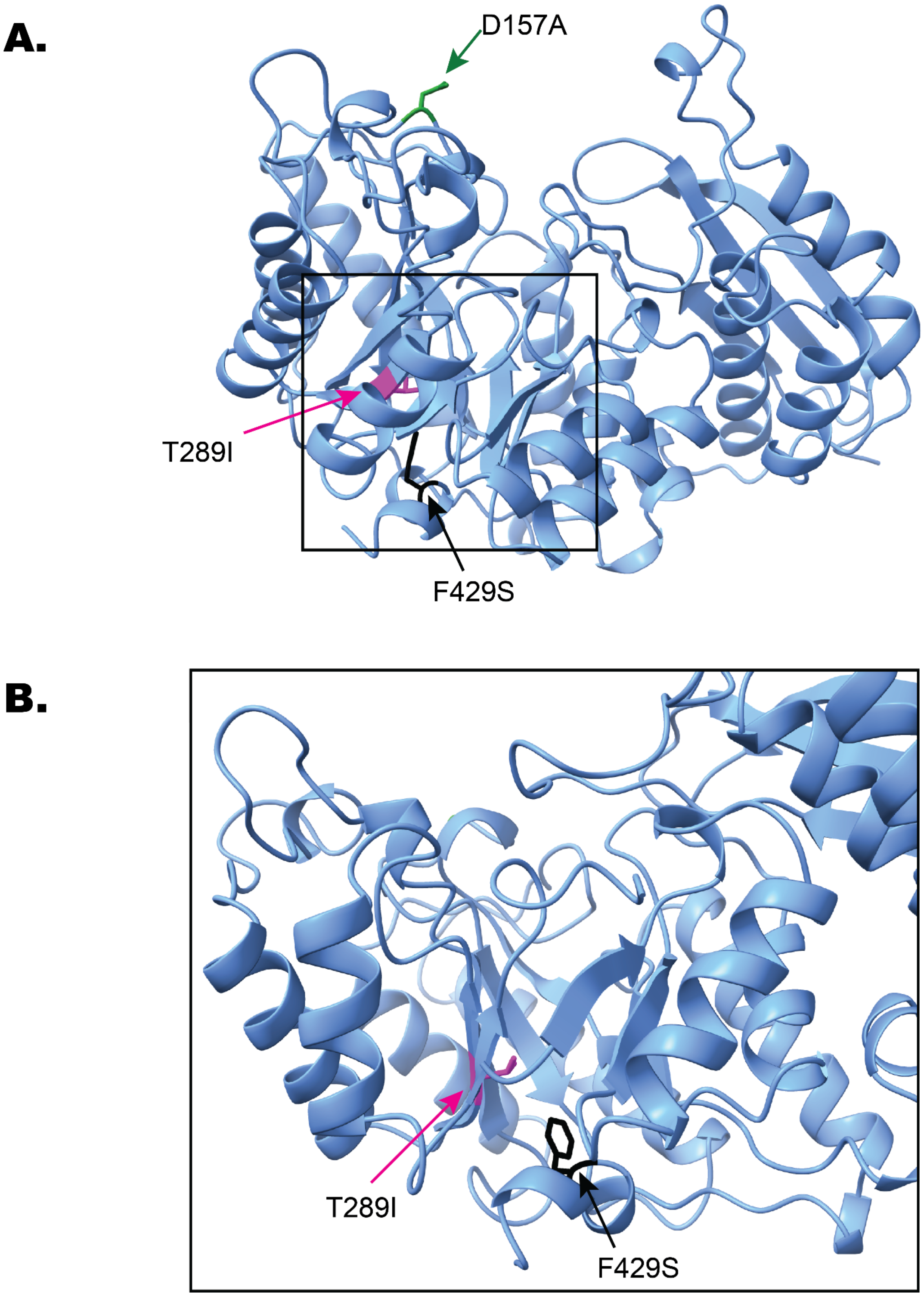

**Supplemental Figure 8:** (**A**) AlphaFold Protein Structure Database Entry AF-A8AY46-F1-v6 for *S. gordonii*'s enolase, sgo_1426. WT positions of mutated residues are marked in green for D157A, magenta for T289I, and black for F429S. Note that this structure does not indicate the predicted conformational changes of the point mutations found in the evolved isolates. Since two of the mutations occurred in internal locations near the active site, panel (**B**) is a zoomed-in look at the WT apo active site from a different perspective, with the sites of the identified mutations highlighted as previously described.

**Supplemental Table 6:** Mutations in enolase, sgo_1426, found in *S. gordonii* isolates sent for WGS. Note that the isolate name indicates the evolutionary condition followed by the population the strain was isolated from.

| Mutation | Isolate |
|---|---|
| T289I | Glucose + Fluoride isolate 1<br>Glucose + Fluoride isolate 2<br>Glucose + Fluoride isolate 3<br>Erythritol + Fluoride isolate 3<br>Sorbitol + Fluoride isolate 1<br>Sorbitol + Fluoride isolate 2<br>Xylitol + Fluoride isolate 3 |
| D157A | Erythritol + Fluoride Isolate 1 |
| F429S | Xylitol + Fluoride Isolate 1 |

**Supplemental Table 7:** Mutation category counts for *S. gordoniii* evolved isolates, with 3 sequenced isolates compiled per evolution environment. Mutation counts are based on the whole genome sequencing processing pipeline described in the methods section.

| Evolution Environment | All non-synonymous mutations | Snp nonsynonymous | Large deletion | Snp intergenic | Snp nonsense | Small indel |
|---|---|---|---|---|---|---|
| Glucose, F | 20 | 15 | 0 | 3 | 0 | 2 |
| Erythritol, F | 12 | 9 | 1 | 1 | 1 | 0 |
| Sorbitol, F | 13 | 10 | 1 | 1 | 0 | 0 |
| Xylitol, F | 18 | 15 | 1 | 2 | 0 | 0 |

**Supplemental Table 8:** Mutation category counts for *S. mutans* evolved isolates, with 3 sequenced isolates compiled per evolution environment. Mutation counts are based on the whole genome sequencing processing pipeline described in the methods section.

| Evolution Environment | All non-synonymous mutations | Snp nonsynonymous | Large deletion | Snp intergenic | Snp nonsense | Small indel |
|---|---|---|---|---|---|---|
| Glucose, F | 10 | 3 | 0 | 4 | 1 | 2 |
| Erythritol, F | 9 | 3 | 0 | 4 | 0 | 2 |
| Sorbitol, F | 10 | 6 | 0 | 3 | 1 | 0 |
| Xylitol, F | 14 | 8 | 0 | 4 | 2 | 0 |

**Supplemental Table 9:** Fluoride $IC_{50}$ of *S. mutans* ancestor and *S. mutans* smu_589 K4Q in varying carbon environments. Data are plotted in Figure 5B.

| Assay Carbon Condition | Strain | mean NaF IC50 (mM) | std_IC50 | IC50_mean ± SD (mM) | Replicates (n) | t-test p-value |
|---|---|---|---|---|---|---|
| 0.1% glucose | *S. mutans* ANC | 4.17 | 0.17 | 4.17 ± 0.17 | 4 | NA |
| 0.1% glucose | *S. mutans* smu_589 K4Q | 4.88 | 0.16 | 4.88 ± 0.16 | 4 | 0.000983 |
| 0.9% erythritol | *S. mutans* ANC | 4.25 | 0.18 | 4.25 ± 0.18 | 4 | NA |
| 0.9% erythritol | *S. mutans* smu_589 K4Q | 5.00 | 0.25 | 5.00 ± 0.25 | 4 | 0.00348 |
| 0.9% sorbitol | *S. mutans* ANC | 1.80 | 0.14 | 1.80 ± 0.14 | 4 | NA |
| 0.9% sorbitol | *S. mutans* smu_589 K4Q | 1.73 | 0.09 | 1.73 ± 0.09 | 4 | 0.445 |
| 0.9% xylitol | *S. mutans* ANC | 4.00 | 0.45 | 4.00 ± 0.45 | 4 | NA |
| 0.9% xylitol | *S. mutans* smu_589 K4Q | 6.36 | 0.31 | 6.36 ± 0.31 | 4 | 0.000247 |

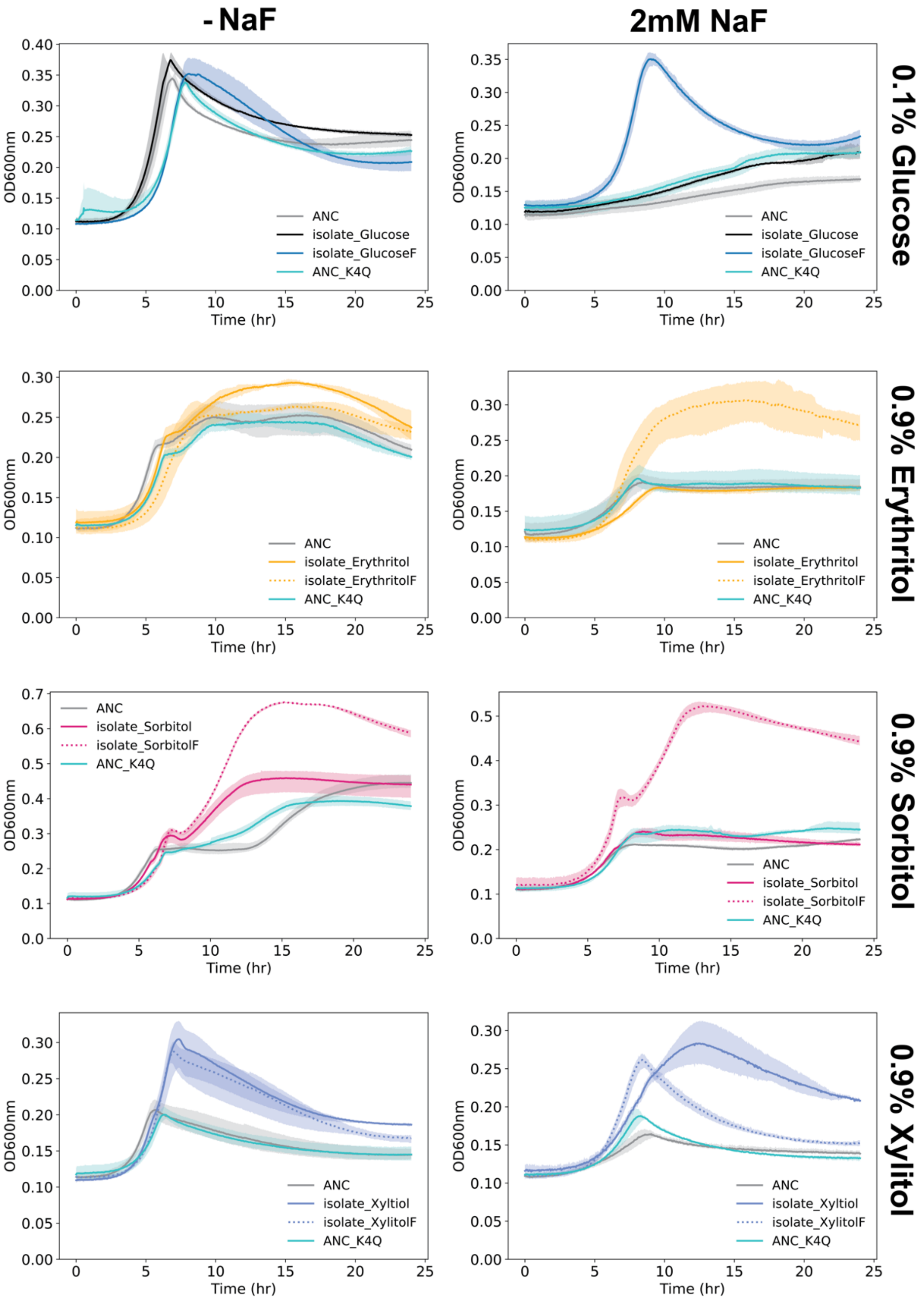

**Supplemental Figure 9:** Growth curves comparing the ancestor (ANC) to the ancestor with the smu_589 K4Q mutation (ANC_K4Q) in the absence (left) and presence (right) of 2mM sodium fluoride in 4 carbon conditions (labeled on the right axis) (n=3). The growth curves of two evolved isolates were included in each plot, one which evolved in only the carbon environment and one that evolved in the carbon environment with fluoride. The isolate randomly selected from population 2 of each condition was used for the growth curve comparison.
